# *Ex Vivo* Culture of Patient-Derived Primary, Metastatic, and Post-Mortem Lung Cancer Reveals Targetable States of EMT and Metabolic Plasticity

**DOI:** 10.64898/2026.06.14.731898

**Authors:** Suehelay Acevedo-Acevedo, Hayley D. Ackerman, Vanessa Y. Rubio, Nicole Hackel, Christina L. Carr, Kaitlyn A. Miranda, Jaden R. Baldwin, Michelle Reiser, John H. Lockhart, Ashley Lui, Paul A. Stewart, Xiaoqing Yu, Gabriela M. Wright, Aileen Y. Alontaga, John M. Koomen, Duy T. Nguyen, W. Gregory Sawyer, Gina M. DeNicola, Theresa Boyle, W. Douglas Cress, Eric B. Haura, Elsa R. Flores

**Affiliations:** Department of Molecular Oncology, H. Lee Moffitt Cancer Center and Research Institute, Tampa, FL 33612, USA; Cancer Biology and Evolution Program, H. Lee Moffitt Cancer Center and Research Institute, Tampa, FL 33612, USA; Department of Nutrition and Integrative Physiology, University of Utah, Salt Lake City, UT 84112, USA; Huntsman Cancer Institute, University of Utah, Salt Lake City, UT 84112, USA; Department of Biostatistics and Bioinformatics, H. Lee Moffitt Cancer Center and Research Institute, Tampa, FL 33612, USA; Department of Pathology, H. Lee Moffitt Cancer Center and Research Institute, Tampa, FL 33612, USA; Department of Bioengineering, H. Lee Moffitt Cancer Center and Research Institute, Tampa, FL 33612, USA; Department of Metabolism and Physiology, H. Lee Moffitt Cancer Center and Research Institute, Tampa, FL 33612, USA; Department of Thoracic Oncology, H. Lee Moffitt Cancer Center and Research Institute, Tampa, FL 33612, USA

**Keywords:** Lung adenocarcinoma, non-small cell lung cancer, small cell lung cancer, *ex vivo* cancer models, tumor progression, epithelial-mesenchymal transition, fatty acid metabolism, rapid tissue donation (RTD), rapid autopsy specimen

## Abstract

Lung cancer is a highly heterogeneous disease and remains the leading cause of cancer-related mortality worldwide. While mouse models and patient-derived organoids have advanced our understanding of lung cancer, key interactions within the tumor microenvironment (TME) remain poorly characterized. We developed microtumor models from lung adenocarcinoma (LUAD) and small cell lung cancer (SCLC) using mouse and patient samples, including surgical resections and rapid autopsy specimens. Microtumors preserve structural, cellular, and molecular features of the native TME, enabling mechanistic studies of tumor progression *ex vivo*. Multi-omics analyses of LUAD microtumors revealed progression-associated changes, including increased epithelial-to-mesenchymal transition (EMT) and metabolic reprogramming toward fatty acid synthesis. Pharmacologic inhibition of fatty acid synthesis through ACC1/2 reduced proliferation in patient-derived microtumors, identifying a targetable vulnerability. This platform provides a robust system for studying tumor progression, therapeutic response, and resistance mechanisms in lung cancer, including culturing postmortem specimens that are not accessible in current models.

## INTRODUCTION

Lung cancer remains the leading cause of cancer-related deaths and the second-leading cancer type contributing to new cases in both males and females^1^. Lung cancer is further divided into two categories: small-cell lung cancer (SCLC) and non-small-cell lung cancer (NSCLC), which encompass 15% and 85% of cases, respectively^2–4^. Among NSCLC, lung adenocarcinoma (LUAD) is the most frequently diagnosed subtype, accounting for approximately 40% of all cases^4,5^. Given the high incidence and mortality of lung cancer, it is essential to develop models that recapitulate the complex tumor microenvironment (TME) to understand tumor initiation, development, metastasis, and response to therapeutics. Over the years, many *in vivo* mouse models of LUAD with different tumor drivers have been developed, including the commonly mutated *Kras* gene^6,7^. While genetically engineered mouse models (GEMMs) provide a means to study LUAD tumor development and progression at the whole-organism level, *in vitro* and *ex vivo* approaches have traditionally been used to gain mechanistic insights into LUAD biology. Similarly, mouse models of SCLC and organoid cultures derived from primary patient tumors or circulating tumor cells have been used to study its subtypes and explore therapeutic vulnerabilities^8–10^. However, experimental systems that enable detailed spatiotemporal analysis of SCLC in the context of dynamic interactions with the TME remain limited.

Culturing lung cancer patient tumors poses significant challenges, such as limited tumor tissue availability and outgrowth of specific cell types in organoid cultures, depending on culture conditions. With advances in bioengineering, many models have been developed to study normal tissue physiology and tumor pathology *ex vivo* in 3D^11^. For example, normal lung tissue from a library of donor tissues was minced and cultured *ex vivo* to evaluate drug response to SARS-CoV-2 infection^12^ and to assess the adherence capacity of different lung cancer cell lines^13^. Additionally, many organ-on-a-chip microfluidics platforms have been developed to study lung cancer^14–16^. These diverse models of lung cancer further highlight the need to recapitulate the endogenous TME to accurately model lung cancer and understand its complex biology. Furthermore, *ex vivo* models of SCLC are limited, as current models rely on isolating circulating tumor cells from liquid biopsies^9,17^ or on short-term culture of patient-derived spheroids^18^. Therefore, it is imperative to establish *ex vivo* models, given the poor patient prognosis and lack of targeted therapies. Here, we developed an *ex vivo* microtumor platform for LUAD and SCLC. This system has previously been used to culture mouse colorectal tumors *ex vivo*, allowing tissues to be cultured in a bioinert microgel scaffold under constant media perfusion for up to 21 days^19^ and to study the response to immunotherapy in a human colorectal cancer sample harboring a pathogenic DNA polymerase ε *(POLE)* mutation.

Building on the success of culturing tumors *ex vivo*, we further leveraged this platform to perform molecular characterization of LUAD and SCLC. Using single-cell sequencing, multi-omics, and immunofluorescence, we analyzed murine-and patient-derived LUAD microtumors cultured *ex vivo* for 10 days under perfusion and compared them with primary tumors and organoids. LuAD microtumors retained epithelial, stromal, and immune cell populations and molecular signatures similar to those of primary tumors, whereas the organoids expressed signatures that diverged from those of the primary tumors. These data indicate that LuAD microtumors recapitulate the biology of the endogenous tumor and the tumor microenvironment. Microtumor cultures of patient specimens derived from LuAD and SCLC samples received from surgical procedures (OR) and the Rapid Tissue Donation (RTD) protocol at Moffitt Cancer Center^21^ were also established using this method. We found that endogenous patient immune cells and markers of tumor identity are preserved within these microtumors, and that we can identify therapeutic vulnerabilities using a multi-omics approach. Furthermore, we demonstrate the platform’s ability to sustain viable patient tissues for an extended period, including those derived from post-mortem RTD samples, thereby providing a novel platform for the analysis of therapy-resistant tumors.

Interestingly, mouse and human LuAD microtumors exhibited upregulated expression of epithelial-to-mesenchymal transition (EMT) genes after *ex vivo* culture. EMT is known to be a critical driver of tumor progression and metastasis^22^ and is heavily influenced by the tumor microenvironment (TME)^23^. Beyond altering the genetic programs in tumor epithelial cells, EMT is known to influence tumor metabolism^24–26^ and is one of the main hallmarks of lipid metabolism reprogramming^27,28^. Importantly, lipidomic analysis of both mouse and human LuAD microtumors revealed lipid reprogramming *ex vivo*. Many studies have aimed to target metabolic vulnerabilities in cancer to disrupt EMT and prevent tumor progression^24^, including the acetyl-CoA carboxylase 1 (ACC1)/ACC2 inhibitor ND-646^29^. Treatment of patient LuAD microtumors with this drug reduced proliferation, demonstrating the strength and scalability of this platform for uncovering new tumor biology and exploiting metabolic vulnerabilities *ex vivo* with ease.

## RESULTS

### Lung adenocarcinomas isolated from mouse models retain key features of the primary tumor after 10 days of *3D* ex vivo culture

*Ex vivo* models that accurately recapitulate LUAD biology are essential for studying tumor progression and developing novel treatment strategies. To address this need, we first developed and characterized 3D *ex vivo* LUAD microtumor models derived from primary mouse LUAD tumors and compared them to the primary tumors and organoids (Figure 1A). All 3 culture models were evaluated by microscopy and multi-omics analysis to characterize the cellular and molecular composition of our *ex vivo* culture.

**Figure 1.**
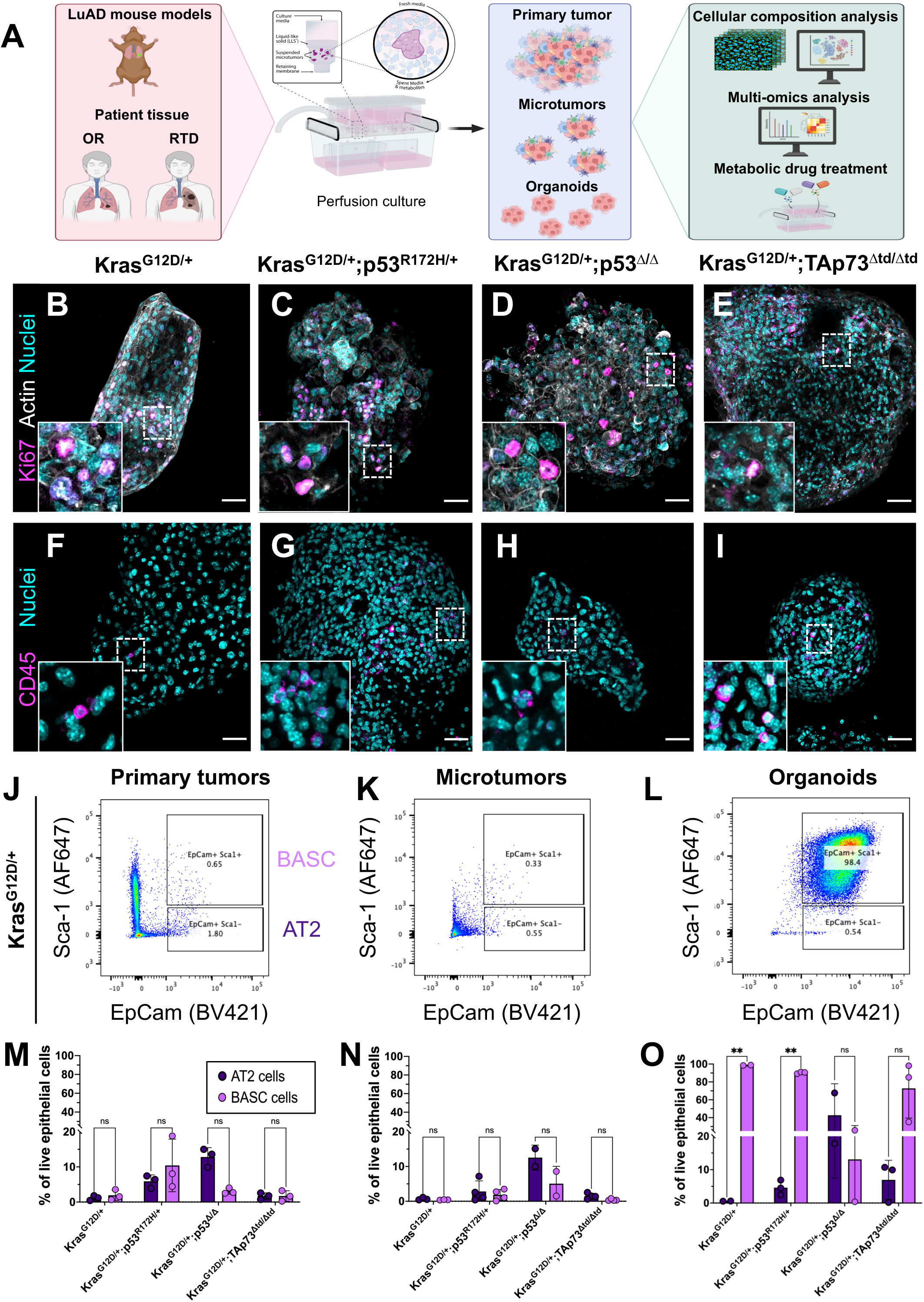
Tumors isolated from mouse models of Kras-driven LUAD retain key features of the primary tumor after 10 days of 3D *ex vivo* culture. A: Schematic detailing the study design using murine and patient-derived samples cultured *ex vivo*. Following culture, the microtumors were characterized at the cellular and molecular level and compared to the primary tumors and organoids. Drug studies were also performed to validate an identified metabolic vulnerability in microtumors. Scheme generated using Biorender. **B – E:** Representative immunofluorescence images of mouse microtumors stained for Ki67 (magenta). Actin is represented in white, and the nuclei in cyan. **F – I:** Representative CD45 (magenta) immunofluorescence images of LUAD microtumors. All scale bars are 50 μm. The dotted white boxes indicate the areas highlighted in the zoomed-in inset images. **J – L:** Representative flow cytometry dot plots of EpCam and Sca1 markers for primary tumor, microtumors, and organoids for the Kras^G12D/+^ mouse model. The upper gates indicate BASC populations, while the lower gates show AT2 cells. **M – O:** Quantification of AT2 and BASC cell populations in primary tumors (M), microtumors (N), and organoid models (O) from *Kras^G12D^*^/+^, *Kras^G12D^*^/+^;*p53^R172H^*^/+^, *Kras^G12D^*^/+^;*p53*^Δ/Δ^, and *Kras^G12D^*^/+^;*TAp73*^Δtd/Δtd^ GEMMs. n = 2–4 mice; multiple pooled tumors per condition. Data shown as mean ± SD. Paired t-tests with Holm-Sidak multiple comparison test were performed using GraphPad Prism (ver 10.6.1). **p < 0.005; ns = not significant.

Generation of these microtumors consisted of harvesting primary tumors from genetically engineered mouse models (GEMMs) and mincing them with surgical scissors into smaller tumors (<1 mm^3^). For *ex vivo* culture, the microtumors were suspended in a bioinert liquid-like solid (LLS) microgel and cultured in commercially available Darcy plates for 10 days under constant media perfusion to maintain tissue viability. At the time of sample preparation for culture, some tissue from the initial tumors was collected for cellular and molecular analysis, while the rest was cryopreserved for future culturing, providing additional versatility to the platform. Furthermore, this platform enables preclinical *ex vivo* testing of novel tumor therapies, including the exploitation of metabolic vulnerabilities in LuAD.

For mouse LuAD models, tumors were collected and cultured *ex vivo* from 4 different LUAD GEMMs: *Kras^G12D^*^/+^, *Kras^G12D^*^/+^;*p53^R172H^*^/+^, *Kras^G12D^*^/+^;*p53*^Δ/Δ^, and *Kras^G12D^*^/+^;*TAp73*^Δtd/Δtd^. To minimize effects arising from differences in culture conditions between the microtumors and organoids, both models were cultured in human plasma-like media (HPLM) with the same nutrient additives. Mouse LUAD microtumors exhibited heterogeneous 3D tumor morphology and had defined actin networks typical of epithelial tumors (Figures 1B–1E). Ki67 immunofluorescence indicated that these microtumors remained proliferative after 10 days in culture (Figures 1B-1E). The microtumors also preserved the endogenous immune microenvironment of the primary tumors, as shown by CD45 staining (Figures 1F-1I).

We also assessed LUAD microtumors for the presence of Alveolar type 2 (AT2) and bronchoalveolar stem cells (BASC), the cells of origin of LUAD tumors^30,31^. Microtumor and organoid samples derived from the GEMMs were assayed for Sca-1 and EpCam positivity by flow cytometry^32^ to determine the proportions of AT2 and BASC cells. Microtumors from Kras^G12D/+^ mice showed that they contained a similar proportion of AT2 (EpCam^+^ Sca1^-^) and BASC (EpCam^+^ Sca1^+^) cells as the primary tumors (Figures 1J-1K), while the organoids showed an expansion of the BASC cell population compared to AT2 cells (Figure 1L). This pattern was also exhibited in microtumors versus organoids derived from *Kras^G12D^*^/+^;*p53^R172H^*^/+^ and *Kras^G12D^*^/+^;*TAp73*^Δtd/Δtd^ mouse LuAD (Figures 1M–1O), while the *Kras^G12D^*^/+^;*p53*^Δ/Δ^ tumors, microtumors, and organoids selected for a larger population of AT2 cells compared to BASC cells. Together, these results demonstrate that LUAD microtumors suspended in the LLS microgels are viable in this perfusion platform and retain endogenous immune cells and stem cell proportions present in the source primary tumors.

### LUAD microtumors recapitulate the cellular and molecular tumor microenvironment

To further characterize the cellular composition of microtumors relative to primary tumors and organoids, we performed single-cell RNA sequencing (scRNA-seq) on samples from *Kras^G12D^*^/+^, *Kras^G12D^*^/+^;*p53^R172H^*^/+^*, Kras^G12D^*^/+^;*p53*^Δ/Δ^, and *Kras^G12D^*^/+^;*TAp73*^Δtd/Δtd^ GEMMs. Approximately 6,000 cells per sample were sequenced and clustered into 13 major cell clusters. The uniform manifold approximation and projections (UMAPs) showed a similar distribution of cell types across primary tumors and microtumors, whereas most cells identified in the organoids grouped within the epithelial cell cluster (Figures 2A-2C). Primary tumors in all GEMMs comprise a heterogeneous mixture of cells characteristic of the tumor microenvironment, including tumor epithelial cells, macrophages, cancer-associated fibroblasts, endothelial cells, and various other immune cell populations, such as B and T cells (Figures 2D-2G). Importantly, the microtumors similarly had a diverse array of cell types, including tumor epithelial cells, fibroblasts, macrophages, endothelial cells, and a smaller percentage of T cells. On the other hand, organoids were highly homogeneous, consisting primarily of tumor epithelial cells, with small proportions of other cell types. These data indicate that organoids selected for tumor epithelial cells, but microtumors maintain a diverse cellular composition similar to the source primary tumors.

**Figure 2.**
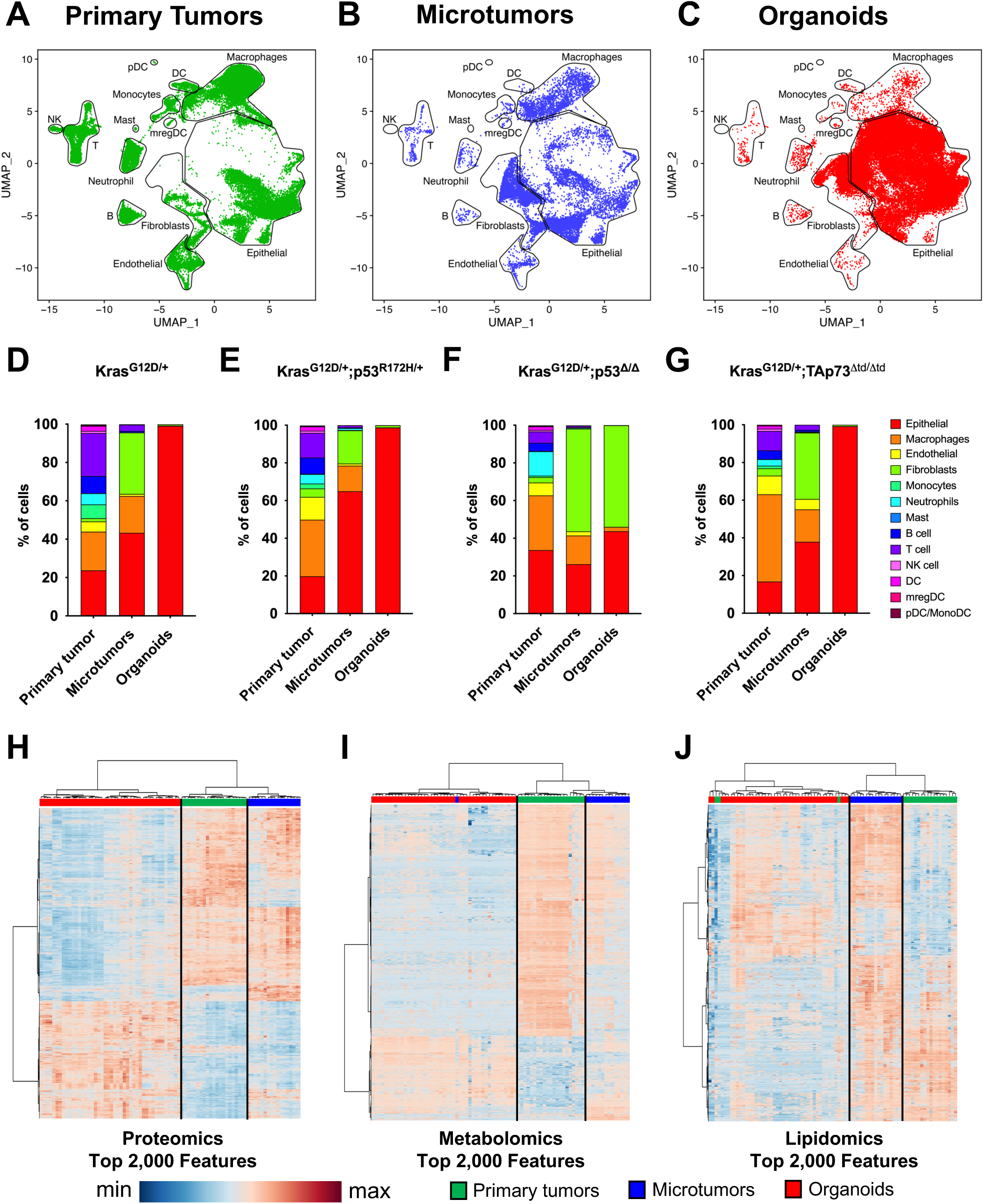
LUAD microtumors recapitulate the cellular and molecular tumor microenvironment in 3D *ex vivo* cultures. A – C: UMAPs of primary tumors (green), microtumors (blue), and organoids (red) for all GEMMs analyzed by single-cell RNA sequencing. **D – G:** Proportion of major cell types for all genotypes. n = 1 – 4 mice per model per genotype, multiple tumors pooled per mouse. **H – J:** Global proteomics, metabolomics, and lipidomics heatmaps of the top 2,000 features for the mouse primary tumor, microtumor, and organoid tissues. Colors indicate z-normalized intensity values. N = 2 – 5 mice per model per genotype, multiple tumors pooled per mouse.

We conducted a broader characterization of molecular differences among primary tumors, *ex vivo* cultured microtumors, and organoids using proteomics, metabolomics, and lipidomics. In all 3 analyses, the microtumors showed a molecular profile similar to that of the primary tumors and clustered closely by unsupervised hierarchical clustering, whereas the organoid molecular profile was discordant with the primary tumors and microtumors (Figure 2H-2J). Taken together, in-depth cellular and molecular characterization shows the microtumors recapitulate the TME and overall molecular profiles of the primary tumors isolated from GEMMs. This suggests that microtumors maintain their tumor identity during *ex vivo* culture, unlike organoids, and recapitulate *in vivo* biological mechanisms that enable them to thrive outside the mouse.

### LuAD microtumors from GEMMs exhibit enhanced EMT signatures in epithelial cells

To identify pathways changed during the extended culturing of microtumors *ex vivo*, we analyzed proteins showing significant differential abundance from the initial primary tumors in the proteomics data. Interestingly, this analysis revealed that the EMT pathway was one of the most enriched in the microtumor samples compared to the primary tumors across all genotypes (Figure 3A). To confirm that this signature was enriched in the malignant epithelial cells, we further analyzed the gene expression in these cells in the scRNAseq analysis of microtumors, primary tumors, and organoids. We evaluated the epithelial clusters against tumor cell state signatures previously reported in *Kras^G12D^*^/+^;*p53*^Δ/Δ^ transplant mice^33–35^ and *Krt8*+ alveolar intermediate cell (KAC) signature^36^. We observed that the primary tumor epithelial cells exhibited gene signatures consistent with previously published malignant epithelial cell signatures^33–35^, such as alveolar type 1-like (AT1-like), AT2-like, gastric-like, and some pre-epithelial mesenchymal transition (EMT) signatures (Figure 3B). Dissimilar from the primary tumors, the organoid epithelial cells demonstrated transition gene signatures, including KACs and pre-EMT expression profiles. Interestingly, the microtumors exhibited high pre-EMT, early EMT, and mesenchymal gene signatures, suggesting that the epithelial cells within the microtumors are progressing toward a more aggressive state. Taken together, this data demonstrates that this 3D platform culture system provides the opportunity to understand and characterize molecular mechanisms in tumor progression that were not previously accessible.

**Figure 3.**
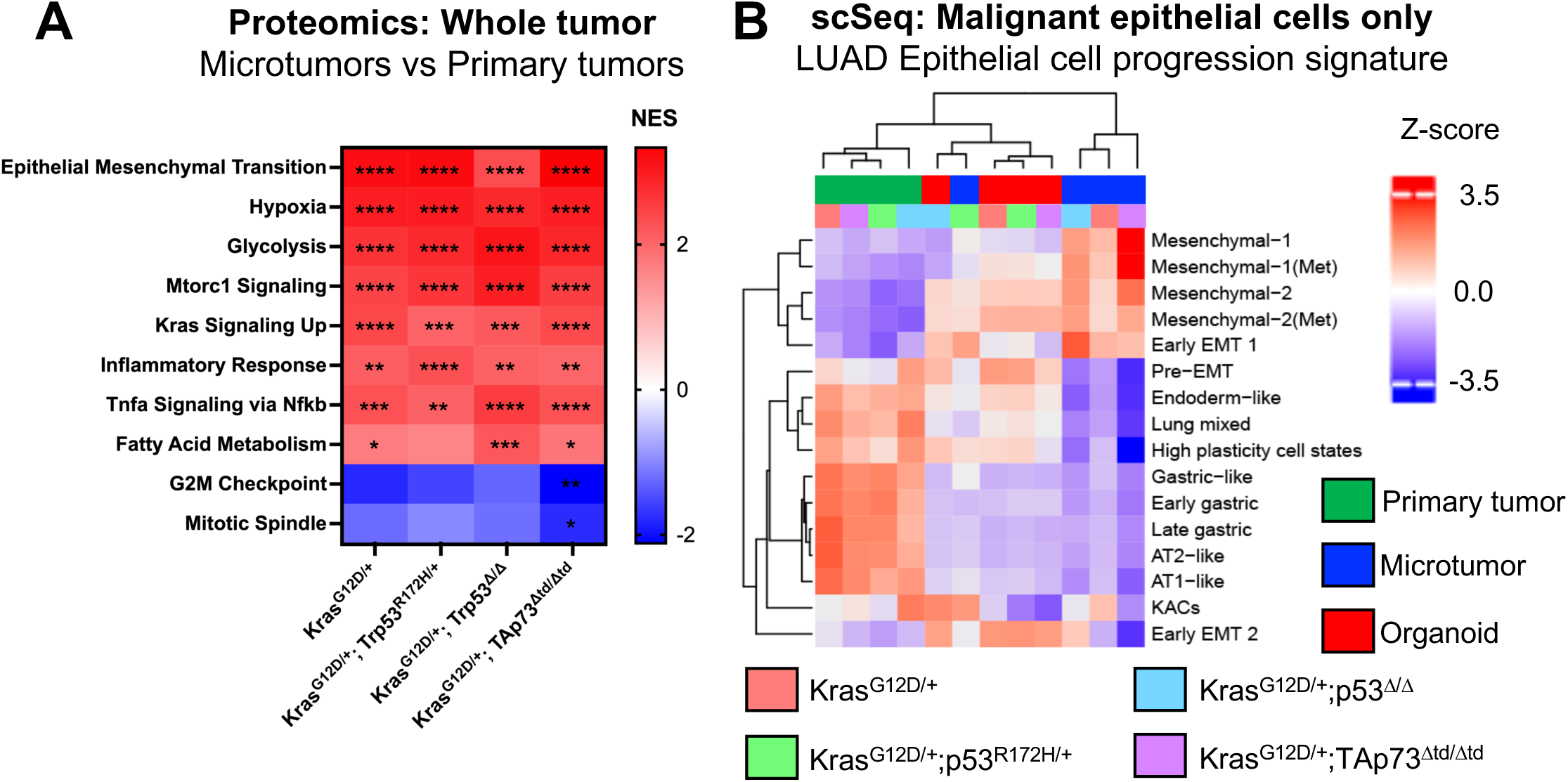
Mouse LUAD microtumors exhibit enhanced EMT signatures in epithelial cells. A: Heatmap showing the combined top proteomic pathways enriched in microtumors compared to primary tumors for all genotypes. Pathways are from the Molecular Signatures Database (MSigDB) hallmark gene set collection^53,54^. Colors indicate the normalized enrichment score (NES). n = 2 – 5 mice per model per genotype, multiple tumors pooled per mouse. FDR q-values calculated by Pre-ranked GSEA^55,56^. *FDR q<0.05; **FDR q<0.005; ***FDR q<0.0005; ****FDR q<0.0001 **B:** Heatmap showing the z-scores of various published gene signatures^33–36^ in malignant epithelial cells from scRNA-seq of the primary tumor (green), microtumors (blue), and organoids (red) for all mouse models. n = 1 – 4 mice per model per genotype, multiple tumors pooled per mouse.

### Lipid metabolism signatures are differentially regulated in lung adenocarcinoma microtumors

EMT is a critical process in the progression of most cancers, including LUAD. Previous research established that lipid metabolism is altered during EMT^27,37,38^, including the accumulation of lipid classes such as sphingolipids^39^ and triglycerides^40,41^. We further examined our lipidomics data to determine whether the microtumors showed lipid reprogramming consistent with LUAD tumor progression. Analysis of global lipidomics confirmed that the microtumors have a distinct lipid profile despite their strong correlation with the primary tumors (Figure 2J). Next, we analyzed the proportions of identified lipid classes to determine which class(es) were modulated by the observed lipid reprogramming (Figure 4A-D). We observed a significant increase in microtumor triglycerides (TG) compared with organoids and primary tumors for all GEMMs. Additionally, ceramides (Cer) were significantly higher for Kras^G12D/+;^TAp73^Δtd/Δtd^ (Figure 4D). Murine LUAD microtumors also had significantly lower levels of phospholipids, including phosphatidylcholine (PC), phosphatidylethanolamine (PE), and phosphatidylglycerol (PG), compared to the primary tumors.

**Figure 4.**
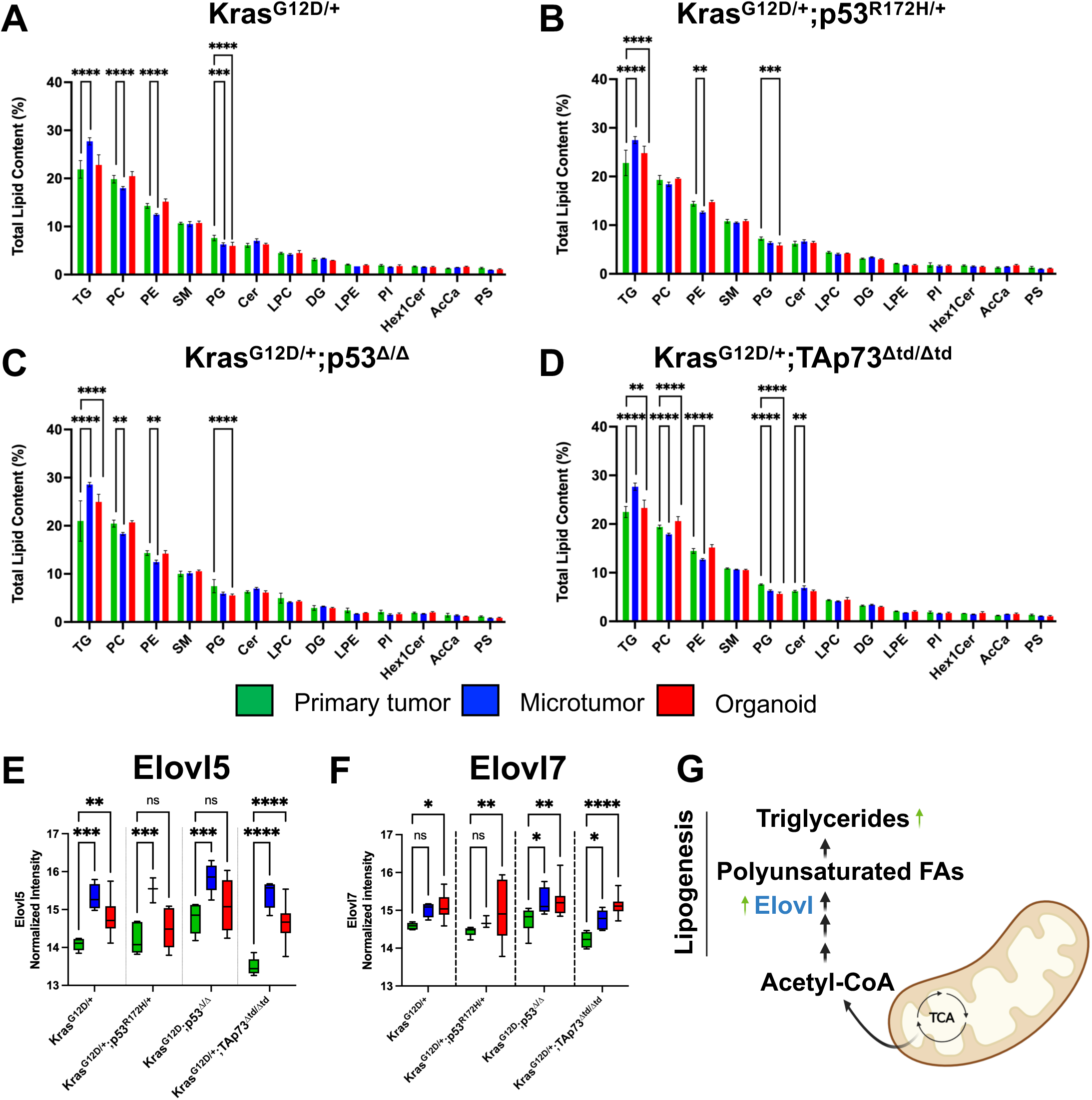
Lipid metabolism signatures are differentially regulated in LUAD microtumors. A – D: Total lipid content percentage for lipid classes identified by lipidomics using Lipid Maps^57,58^ in primary tumors, microtumors, and organoids for all four GEMMs. TG = Triglyceride, PC = Phosphatidylcholine, PE = Phosphatidylethanolamine, SM = Sphingomyelin, PG = Phosphatidylglycerol, Cer = Ceramide, LPC = Lysophosphatidylcholine, DG = Diglyceride, LPE = Lysophosphatidylethanolamine, PI = Phosphatidylinositol, Hex1Cer = Hexosylceramide, AcCa = Acylcarnitine, PS = Phosphatidylserine. Multiple unpaired t-tests with a single pooled variance and Holm-Sidak’s multiple comparisons test were performed. n = 4-12 replicates per culture condition. **E – F:** Normalized intensity values for Elovl5 and Elovl7 proteins in the proteomics analysis of all GEMMs. Data shown as min to max box plots. An ordinary two-way ANOVA with uncorrected Fisher’s LSD, with a single pooled variance, was performed. n = 4-12 replicates per genotype per culture condition. All statistical analyses were performed in GraphPad Prism (ver. 10.6.1). *p<0.05; **p<0.005; ***p < 0.0005; ****p < 0.0001; ns = not significant. **G:** Schematic summarizing the observed changes in lipid metabolism during microtumor culture. Generated using Biorender.

Interestingly, the fatty acid metabolism pathway was also significantly enriched in the microtumor compared to primary tumors in proteomics data (Figure 3A). Given elevated triglyceride levels in the microtumor lipidome, we examined the expression of lipid-synthesis enzymes in our proteomics data. Key fatty acid metabolism enzymes Elovl5 and Elovl7 (Figure 4E and 4F) were increased in microtumors compared to the primary tumors, supporting increased lipogenesis and triglyceride synthesis in these tumors (Figure 4G). These results are consistent with previously reported changes in EMT and tumor progression and further emphasize the utility of this 3D *ex vivo* platform as a tool to study novel pathways driving tumor progression.

### LUAD and SCLC microtumors from patient OR and rapid tissue donation (RTD) sources are viable ex vivo

Successfully culturing patient tumors *ex vivo* presents both substantial challenges and significant opportunities. This platform has previously been shown to preserve the viability of patient-derived colorectal microtumors¹⁰. Building on our success in generating microtumors from mouse models (Figures 1 and 2), we next applied this platform to culture lung cancer patient tumors *ex vivo*. We cultured lung adenocarcinoma (LUAD) tumors from 14 operating room (OR) patients as microtumors for 10 days (Table 1). These tumors were largely treatment-naïve and derived from early-stage disease (Table S1). In parallel, we attempted to establish organoid cultures from the same OR patient samples; however, these cultures were predominantly overgrown by normal epithelial cells. Microtumor viability was assessed using live/dead cell staining and fluorescence imaging, with representative images shown in Figures S1A–S1C. Cultures were classified as viable (>80% live cells), moderately viable (20–80% live cells), or non-viable (<20% live cells). After 10 days in culture, 78.6% of OR LUAD microtumors were viable, and 21.4% were moderately viable (Figure S1D). In addition to freshly resected samples, microtumors were successfully established from tumors that had been cryopreserved or stored overnight at 4 °C in HypoThermosol FRS (Tables 1 – 3). Together, these results demonstrate that this platform supports the *ex vivo* expansion of primary patient tumors and enables biobanking for future studies.

**Table 1.**
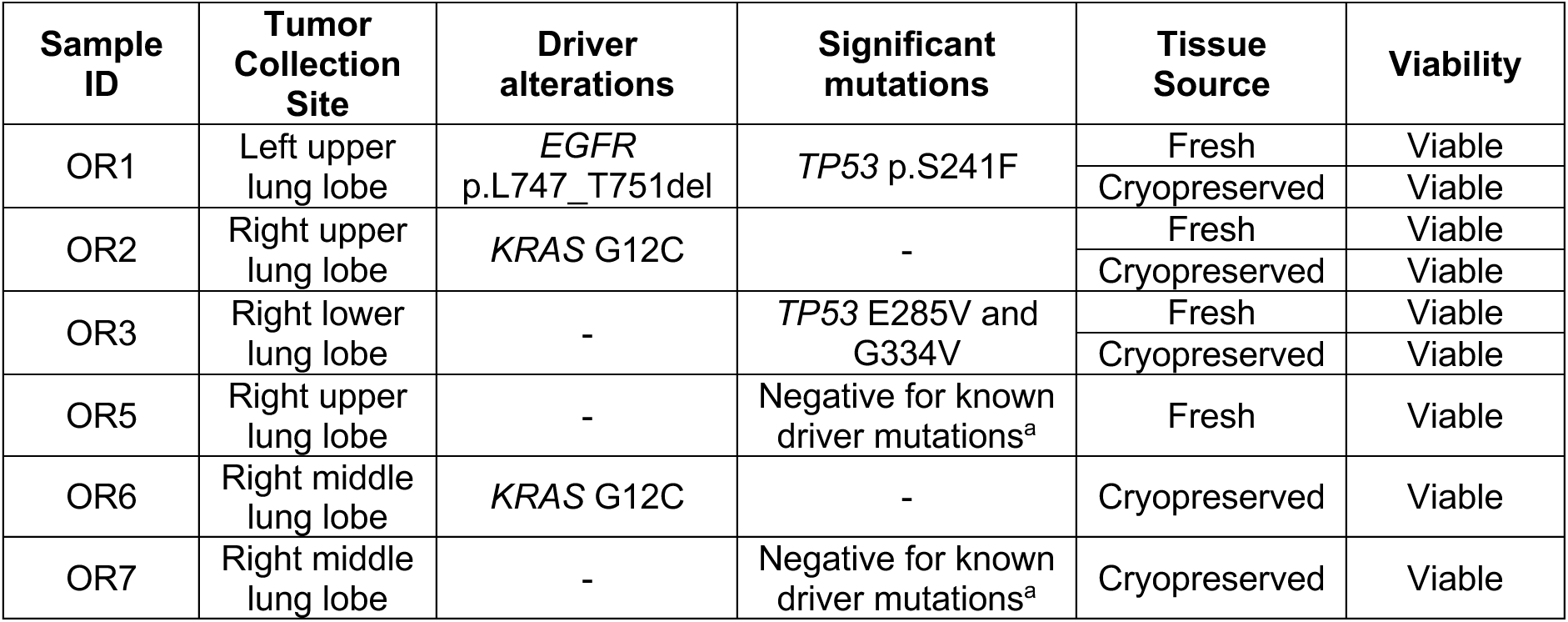
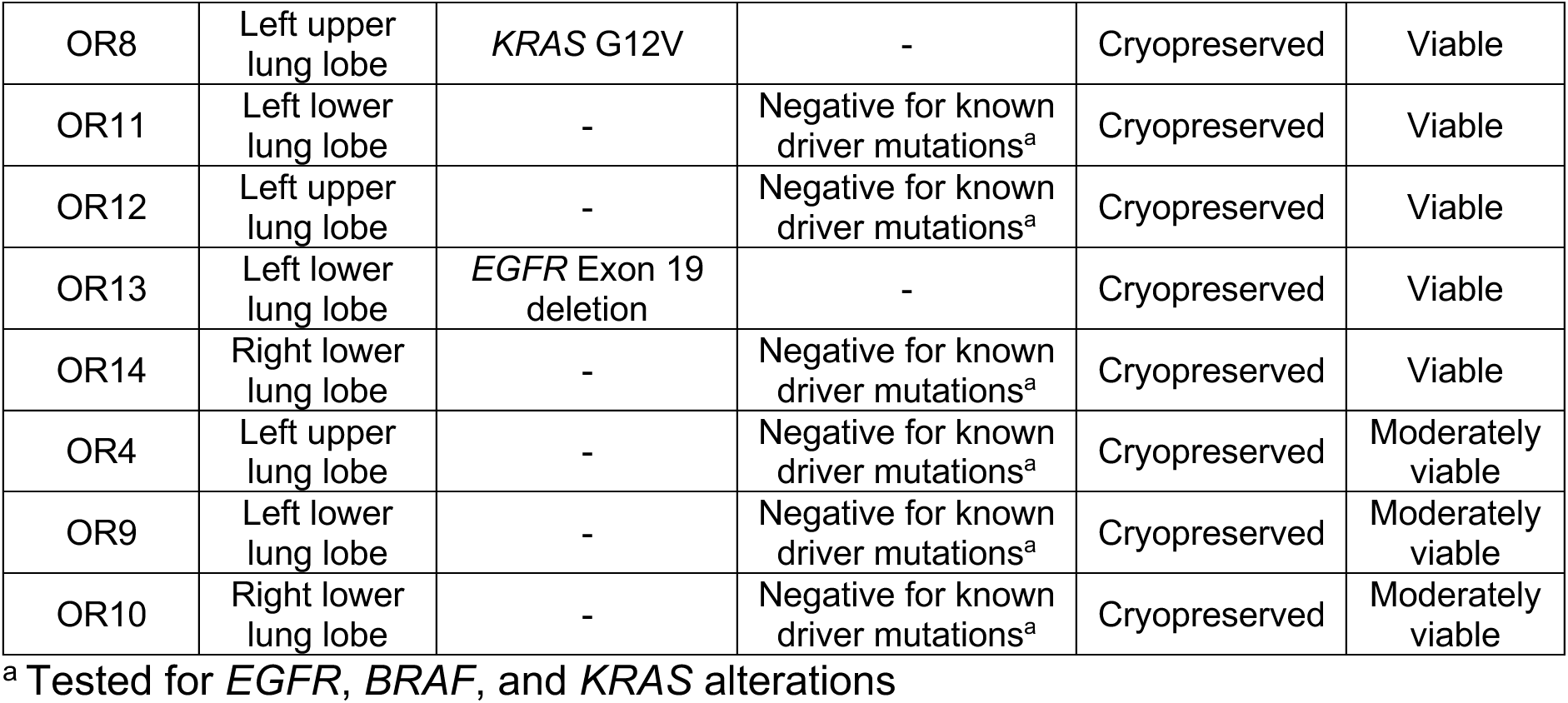
Summary of OR LUAD patient microtumors cultured in 3D.

Building on our success with culturing OR samples in 3D, we asked whether we could also culture and analyze tumors from our rapid tissue donation (RTD) program. To do this, we established *ex vivo* cultures from 11 RTD specimens, including 8 primary LUAD tumors and 3 liver metastases collected from 8 patients through the Rapid Tissue Donation (RTD) protocol^21^ (Table 2). All patients had extensive prior treatment (Table S2), providing a unique opportunity to interrogate tumors that had evolved therapeutic resistance. Notably, all three liver metastasis samples remained viable after 10 days in culture. In contrast, the successful culture of primary lung tumors was highly dependent on ischemic time (Figure S1E). Delayed autopsy leading to ischemic intervals of ≥20 hours markedly compromised viability, yielding 45.5% viable, 27.3% moderately viable, and 27.3% non-viable primary tumor samples (Figure S1E). Despite this sensitivity, we successfully cultured five RTD primary LUAD tumors from fresh or cryopreserved tissue for at least 10 days, demonstrating the feasibility of generating viable *ex vivo* models even from heavily pretreated end-stage disease.

**Table 2.**
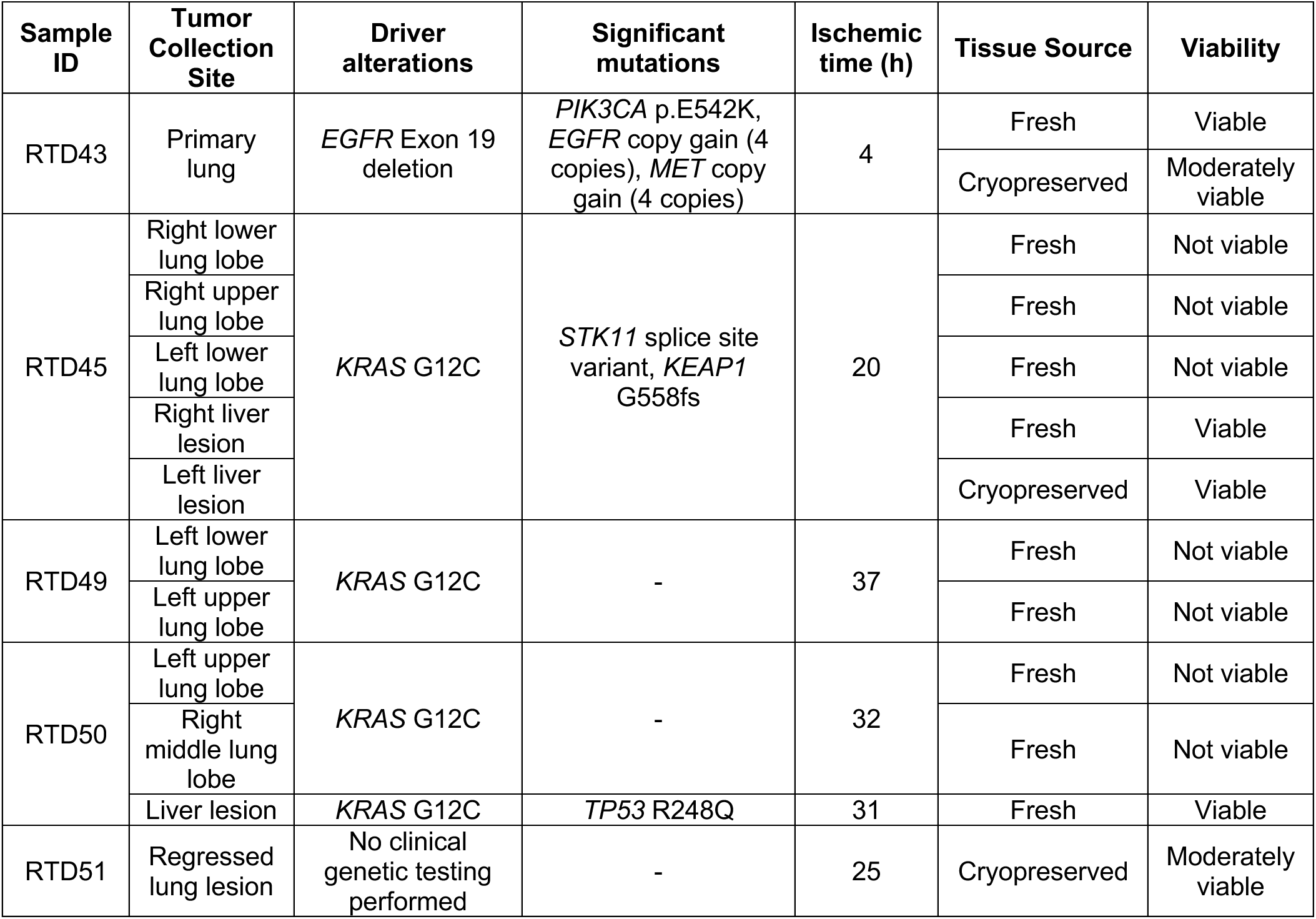
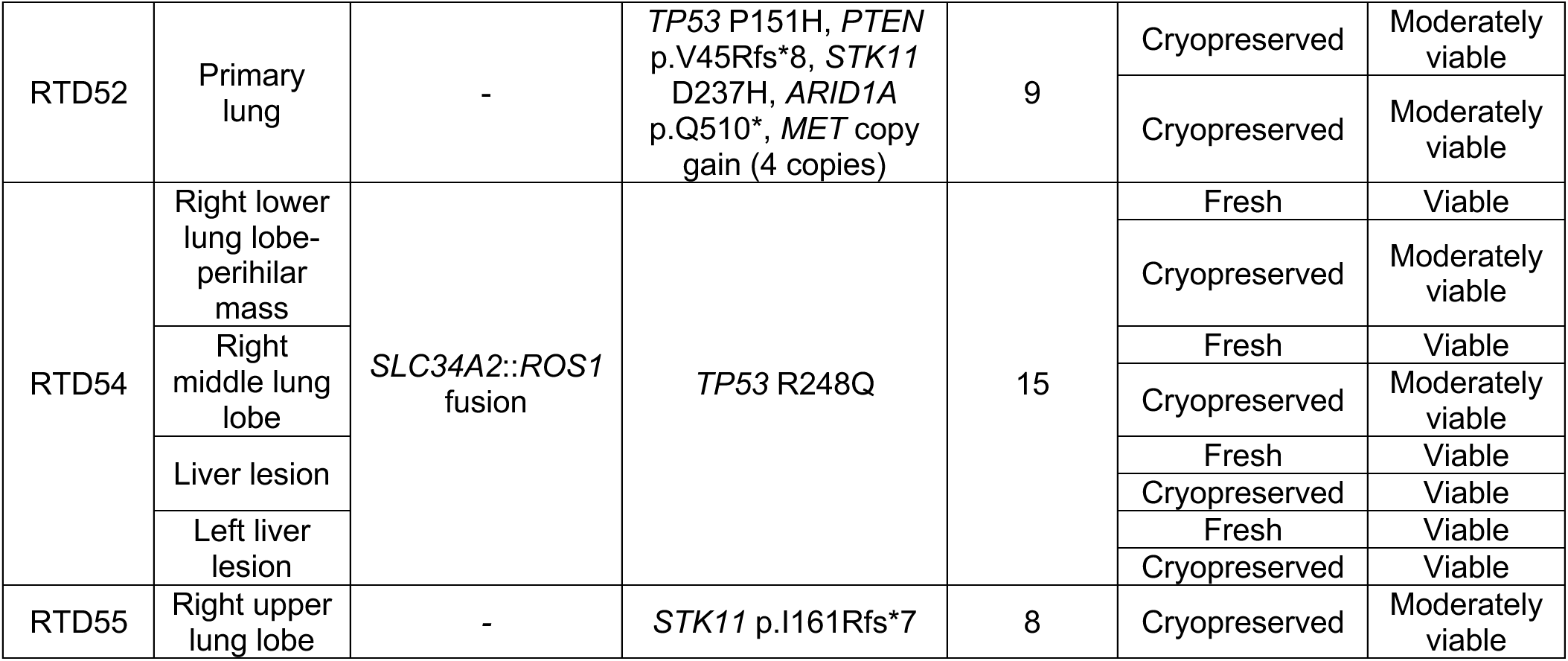
Summary of rapid tissue donation (RTD) LUAD microtumors cultured in 3D.

In addition to LUAD, we established robust *ex vivo* cultures from small cell lung cancer (SCLC) patient samples obtained through the Rapid Tissue Donation (RTD) program, addressing a critical unmet need created by the near absence of surgically resected SCLC specimens. From 10 RTD-derived tumors—6 primary lung cancers and 4 liver or brain metastases collected from six patients—we generated viable cultures representing multiple clinically relevant SCLC subtypes defined by neuroendocrine lineage programs (NEUROD1 or ASCL1), as determined by the custom RNA Salah Targeted Expression Panel (RNA STEP) (Table 3). Notably, 90% of samples remained viable following establishment from fresh or cryopreserved tissue (70% highly viable, 20% moderately viable; Figure S1F), underscoring the reproducibility and scalability of this approach. Primary SCLC lung samples demonstrated greater tolerance to extended ischemic intervals than LUAD RTD tissues (Table 3), greatly expanding the temporal window for tissue acquisition in real-world clinical settings. Collectively, these findings establish a clinically actionable platform for generating patient-representative SCLC models that capture intra-and inter-patient heterogeneity across primary and metastatic sites. To our knowledge, this is the first demonstration that primary and metastatic LUAD and SCLC tumors derived from post-mortem patient tissue can be sustained long-term *ex vivo*, including after cryogenic preservation—creating new opportunities for subtype-specific drug testing, biomarker discovery, and therapeutic development in a disease historically inaccessible to functional precision oncology approaches.

**Table 3.**
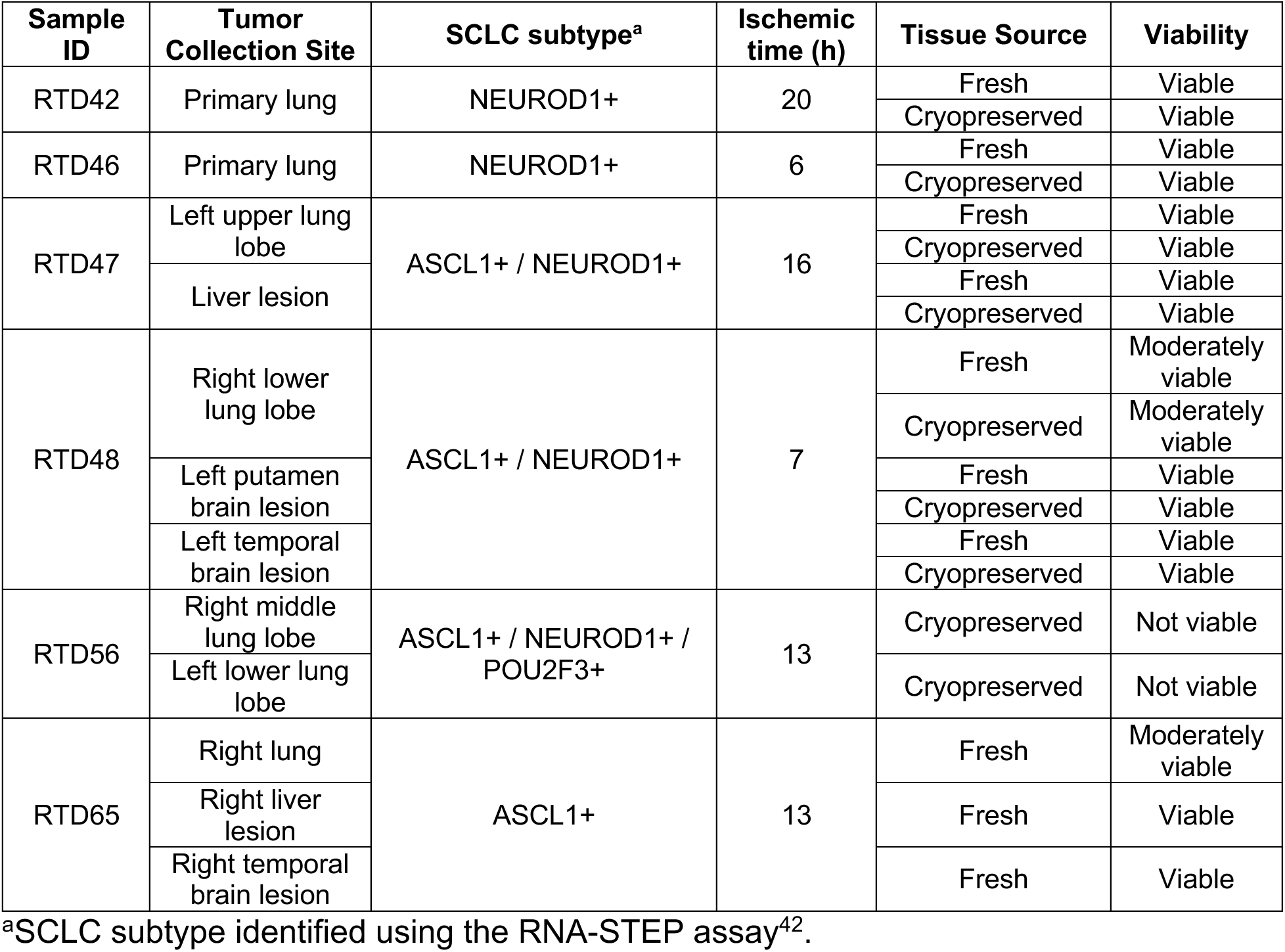
Summary of rapid tissue donation (RTD) SCLC microtumors cultured in 3D.

### Patient-derived lung tumors cultured ex vivo maintain the endogenous tumor microenvironment

To further characterize the patient microtumors, we immunostained the microtumors derived from OR primary lung tumors, RTD-derived primary lung tumors, and RTD-derived metastasis for CD45 to determine if they retained immune cells in the TME. Similar to microtumors derived from GEMMs, the microtumors from LUAD and SCLC patients preserved the endogenous patient immune cells, as evidenced by the presence of CD45-positive cells after 10 days of *ex vivo* culture (Figure 5A-5F).

**Figure 5.**
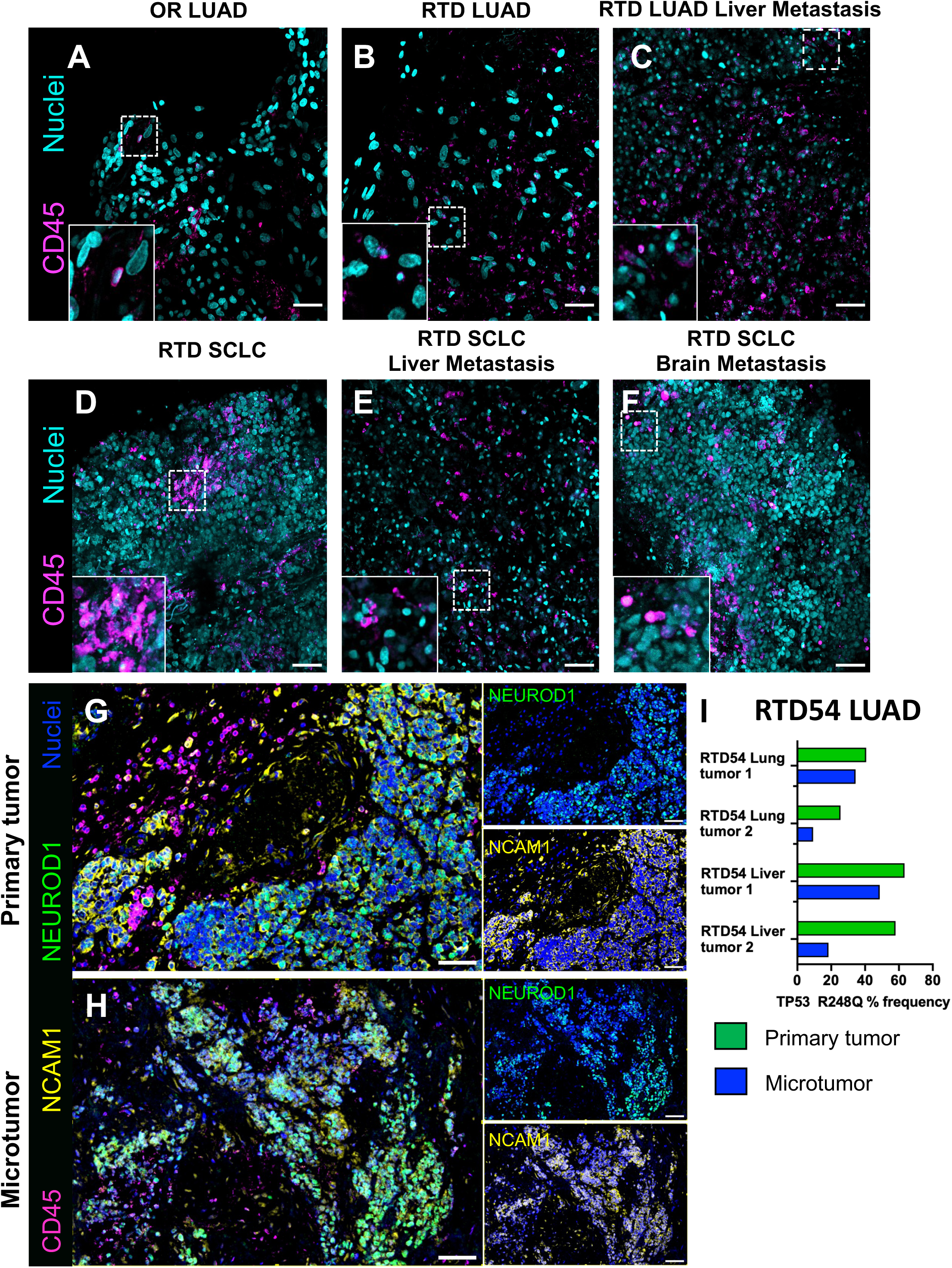
Patient-derived lung tumors cultured *ex vivo* maintain the endogenous tumor microenvironment. A – F: Representative CD45 (magenta) immunofluorescence images of patient microtumors from OR, RTD, or RTD metastatic sites for LUAD and SCLC tumors. Nuclei are shown in cyan. Scale bar = 50 µm. The dotted white boxes indicate the areas highlighted in the zoomed-in inset images. **G – H:** Representative multiplex immunofluorescence images showing NCAM1 (yellow) and NEUROD1 (green) staining for RTD42 SCLC primary lung tumor and a corresponding microtumor after 10 days of *ex vivo* culture. Immune cells (CD45+) are shown in magenta, and nuclei are in blue. Single-color images for NCAM1 and NEUROD1 are shown on the right side of the merged images. Scale bar = 100 µm. **I:** Graph showing the frequency of *TP53* R248Q mutation present in the RTD54 patient sample initially and after microtumor culture for primary lung tumors and liver metastasis sites. Mutation frequency determined via the Moffitt STAR 2.0 panel^43,59^.

Next, we wanted to evaluate if the microtumors retained specific tumor markers and mutations from the primary tumors. For SCLC patient samples, we evaluated whether the microtumors expressed the common SCLC tumor marker NCAM1 (CD56) and retained the same subtype markers identified by RNA Salah Targeted Expression Panel (STEP) analysis^42^ (Table 3). For example, we assayed sample RTD42 and observed that the microtumors were NCAM1 positive after 10 days of *ex vivo* culture and expressed NEUROD1, similar to the primary tumor (Figure 5G-5H).

For the LUAD samples, genetic testing of the microtumors using the Moffitt STAR 2.0 panel^43^ (an RNA-and DNA-based next-generation sequencing assay that assesses small variants and alterations in 500 genes) was performed to ensure that any existing mutations in patients were accurately identified and preserved. For example, genetic testing of the patient sample RTD54, which harbored a *TP53 R248Q* mutation in both primary lung tumors and metastatic liver lesions, showed that all tissues retained the mutation after *ex vivo* culture (Figure 5I).

### Human LUAD microtumors exhibit markers of tumor progression in 3D ex vivo culture

We next wanted to evaluate whether the OR patient-derived LUAD microtumors also exhibited the EMT signatures and lipid metabolic reprogramming that we had observed in the LUAD microtumors from GEMMs. Quantitative PCR was performed to assess the expression of epithelial genes *CDH1* (E-cadherin) and *MUC1* and mesenchymal genes *AXL* and *VIM* (vimentin) in primary tumors and microtumors (Figure 6A). Interestingly, microtumors consistently upregulated both epithelial and mesenchymal genes relative to primary tumors, suggesting a transitional EMT state, as described in more aggressive cancer cell types^44,45^.

**Figure 6.**
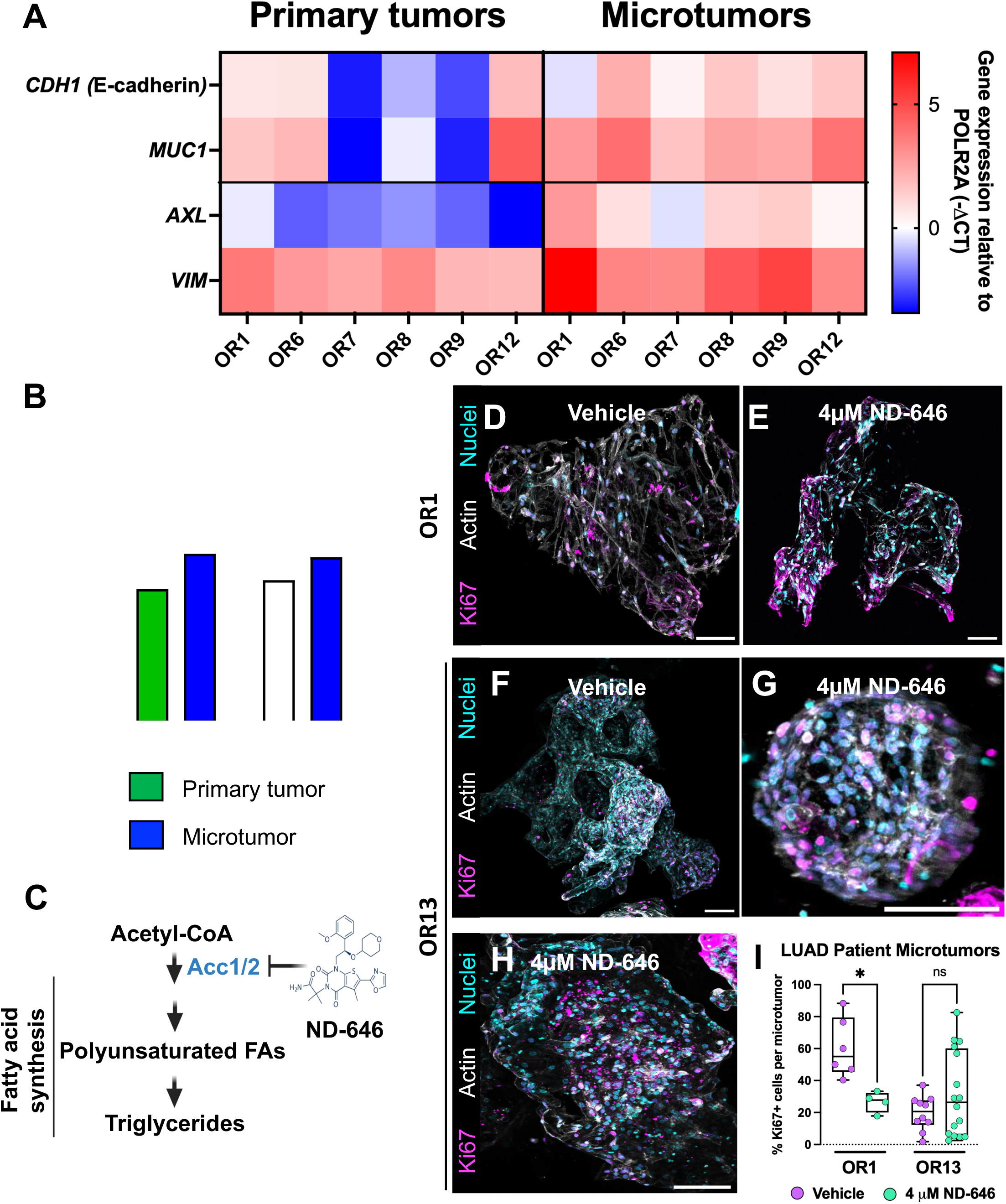
LUAD microtumors exhibit markers of tumor progression and are sensitive to lipid synthesis inhibition. A: Heatmap showing the relative expression levels of genes involved in EMT for the primary and microtumors of lung OR patient samples. Gene expression was measured by qPCR and normalized to POLR2A. **B:** Quantification of the percentage of total lipid content for triglycerides for OR and RTD patient tumors and microtumors. n = 3 – 5 patient samples per condition. Data shown as mean ± SD. An unpaired t-test with Holm-Sidak’s multiple comparisons test was performed using GraphPad Prism (ver. 10.6.1). *p<0.05; ns = not significant. **C**: Diagram showing the target enzymes inhibited by ND-646 within the lipid synthesis pathway. Generated using Biorender. **D – H:** Representative Ki67 (magenta) images of OR microtumors from patients OR1 and OR13 treated with 4 µM ND-646 or Vehicle for 96 hours. Nuclei are shown in cyan, and actin in white. Scale bars: 100 µm. **I:** Quantification of Ki67+ cells in microtumors from OR patients treated with vehicle or 4 µM ND-646 for 96 hours. n = 4 – 6 microtumors per condition. An unpaired t-test was performed using GraphPad Prism (ver 10.6.1). **p<0.005.

Analysis of untargeted lipidomic data revealed no difference in triglyceride levels between primary tumors and cultured microtumors from OR patients (Figure 6B). However, RTD lung microtumor samples showed a significant increase in triglyceride levels compared with primary tumors (Figure 6B). This change suggests that more advanced tumors, such as the RTD samples, which are typically at later stages of tumor progression, exhibit lipid reprogramming signatures similar to those observed in our mouse microtumors. Interestingly, the OR samples did show a trend towards higher triglycerides, suggesting that with time, these tumors could show stronger signals of progression. Taken together, this data supports that the microtumor platform enables mechanisms of tumor progression, such as the upregulation of EMT markers and lipid reprogramming.

### Lipid synthesis was identified as a therapeutic vulnerability in lung adenocarcinoma

One advantage of culturing these microtumors *ex vivo* is the ease with which new therapeutic targets can be identified and assessed, especially in advanced tumors. Based on the observed lipid reprogramming in microtumors that favors lipid synthesis, we treated patient microtumors with ND-646, a small-molecule inhibitor of key lipid biosynthesis enzymes Acc1 and Acc2^29,46^ (Figure 6C). This drug has shown some success in reducing tumor volumes in mouse models of non-small cell lung cancer and could be a promising drug to target metabolic vulnerabilities that arise during tumor progression. Two OR patient lung adenocarcinomas (OR1 and OR13) were treated with either vehicle or 4 µM ND-646 for 96 hours, and cell proliferation was quantified by Ki67 fluorescence imaging. Interestingly, in patient OR1 microtumors, ND-646 treatment reduced cell proliferation, as shown by a decrease in Ki67-positive cells (Figures 6D-E and 6I). For patient OR13, we observed two distinct populations: some microtumors responded to drug treatment, while others remained resistant (Figures 6F-I). These data suggest that patient-derived microtumors can be used to assay new drug therapeutics at early stages of tumor progression while capturing the inherent tumor heterogeneity observed in patients.

Overall, the data demonstrate the establishment of an *ex vivo* lung cancer microtumor culture platform that recapitulates the tumor microenvironment and the global molecular signatures of primary tumors in both mouse and human models. Likewise, it preserves tumor viability for over 10 days with minimal processing and supports the survival of difficult-to-culture samples, such as LUAD and SCLC patient RTD tumors. Furthermore, this platform enables additional modeling of intermediate LUAD tumor progression, EMT, and lipid-reprogramming signatures *ex vivo*. Microtumors can also serve as a powerful tool for screening drugs that modulate novel therapeutic targets.

In summary, we comprehensively characterized murine microtumors and demonstrated that they faithfully recapitulate the cellular and molecular features of their parent tumors. We developed a robust *ex vivo* culture system using patient-derived LUAD and SCLC microtumor samples from both primary and metastatic sites, including difficult-to-access post-mortem RTD specimens. Importantly, this platform captured key hallmarks of tumor progression, including EMT and lipid metabolic rewiring, thereby enabling mechanistic interrogation of disease evolution and exposing a potential therapeutic vulnerability.

## DISCUSSION

Lung cancer is a heterogeneous disease in which the TME contributes to tumor progression, metastasis, and therapeutic response and resistance. Therefore, it is essential to generate *ex vivo* models that retain the TME and mimic the biology of this complex disease. We established *ex vivo* models of LUAD from mouse and human tumor samples and demonstrated their utility for studying early tumor progression and metabolic reprogramming. Additionally, we established *ex vivo* SCLC models from post-mortem rapid tissue donation (RTD) patient samples. The generation and culture of lung cancer microtumors in these pressure-gradient-driven perfusion plates enabled a significant extension of *ex vivo* culture duration, as has also been shown with mouse and patient colorectal microtumors^19,20^.

We first used mouse LUAD tumors from diverse genotypes for extensive cellular and molecular characterization of these microtumors. A multi-omics approach showed a strong correlation of the microtumors with the primary tumors from which they were derived (Figures 1 and 2). Therefore, mouse microtumors can mimic the complexity of LUAD biology *ex vivo* and can be studied in a more mechanistic manner. Interestingly, mouse and human LUAD microtumors exhibited upregulated expression of epithelial-to-mesenchymal transition (EMT) genes after *ex vivo* culture. EMT is known to be a critical driver of tumor progression and metastasis^22^ and is heavily influenced by the TME^23^. Understanding EMT biology is crucial to developing novel therapeutics that can target this process. We observed that both human and mouse microtumors upregulated EMT-related genetic and protein markers compared with primary tumors and organoids (Figures 3 and 6). Previous studies have replicated aspects of EMT *in vitro* using microfluidic devices or hydrogel scaffolds, and by adding EMT-promoting factors to the cell culture medium. However, the advantage of using microtumors to study LUAD disease progression and EMT is that the endogenous tumor microenvironment in mouse and human tissues is maintained.

Beyond altering the genetic programs in tumor epithelial cells, EMT is known to influence tumor metabolism^24–26^, and one of its main hallmarks is lipid metabolism reprogramming^27,28^. We observed an increase in triglyceride synthesis pathways in microtumors, which correlates with EMT progression (Figures 4 and 6). Many studies have aimed to target metabolic vulnerabilities in cancer to disrupt EMT and prevent tumor progression^24^, including treatment with the acetyl-CoA carboxylase 1 (ACC1)/ACC2 inhibitor ND-646, previously shown to reduce tumor volume in LUAD cell lines and in *Kras^G12D^;p53^−/−^*and *Kras^G12D^;Stk11^−/−^* mouse models^29^, to block lipid synthesis. We observed reduced cell proliferation in patient OR1 treated with the ND-646 inhibitor after 96 hours (Figure 6), demonstrating the strength of this platform for uncovering new tumor biology and exploiting metabolic vulnerabilities *ex vivo* with ease.

Interestingly, patient sample OR13 showed greater variability in the proliferation response, underscoring the inherent heterogeneity of tumor responses to therapies. These differences could be due in part to the genetic makeup of the patients’ tumors. While both patient tumors had EGFR deletions and similar tumor staging, patient OR1 also had a p53 mutation that could have contributed to the sensitivity of the microtumors to ND-646 treatment. Additionally, the process of generating microtumors from the primary tumors encompasses different tumor regions, including those that are more resistant to treatment. Therefore, using this platform to screen potential therapies in resected tumors could enable the identification of resistance mechanisms and more accurate predictions of the most effective treatments.

Patient-derived models include establishing cell lines, organoids, xenografts, microfluidics, and tissue slices, among others^49^. This *ex vivo* culture method supports the viability, TME, and biology of samples from multiple sources, including surgical OR tissue and post-mortem RTD tissue, due to the minimal tissue processing prior to culture (Tables 1-3, Figure S1, Figure 5). This methodology also allows tissues to be cryopreserved and cultured later, enabling biobanking and extended study and testing of patient samples. Importantly, our successful culture of RTD patient tissues provides a significant advantage for this model in studying mechanisms of tumor progression, metastasis, and even drug resistance in advanced LUAD and SCLC. These microtumors are an essential tool for further understanding these cancers, particularly SCLC, for which few *ex vivo* patient-derived models have been described in the literature.

Collectively, these findings establish the *ex vivo* lung cancer microtumor platform as a rigorously validated, physiologically faithful system that closes a critical gap between primary tumors, metastasis, and experimental models. Its inherent adaptability positions microtumors as a scalable and powerful platform for functional interrogation of therapeutic vulnerabilities and drug response, with strong and immediate relevance for translational research and precision oncology.

## RESOURCE AVAILABILITY

### Lead contact

Requests for further information and resources should be directed to and will be fulfilled by the lead contact, Dr. Elsa R. Flores.

### Materials availability

All materials used in this study, including mouse models, are commercially available unless stated otherwise. *The Kras^G12D^*^/+^;*TAp73*^Δtd/Δtd^ mouse line is available on request to the lead contact, but we may require payment and/or a completed materials transfer agreement if there is potential for commercial application.

### Data and code availability

New scRNA-seq data have been prepared for upload to GEO (Accession #: pending). The mass spectrometry proteomics data have been deposited in the ProteomeXchange Consortium via the PRIDEpartner repository (dataset identifier: pending). The mass spectrometry metabolomics and lipidomics data have been deposited in MetaboLights^52^ (dataset identifiers: pending). All datasets are publicly available as of the date of publication.

This paper does not report original code. Any additional information required to reanalyze the data reported in this paper is available from the lead contact upon request.

## ACKNOWLEDGEMENTS

Initial funding for this work was provided by an NIH/NCI R35-CA197452 to E.R.F. Current and ongoing support for this work has been provided by NIH/NCI P01-CA250984 to E.R.F. The funders played no role in the study design, data collection, analysis, interpretation of data, or the writing of this manuscript. The Salah Foundation generously funded the validation and testing of the RNA STEP. We thank Dr. Ben Creelan and Jyllica Aurelio for providing fresh tissue from lung cancer surgery cases (OR samples). The rapid tissue donation (RTD) project was funded by the James and Esther King Biomedical Research Award by the Florida Department of Health, and the Moffitt Lung Cancer Center of Excellence. We sincerely thank the patients and their loved ones for graciously consenting to participate in the thoracic RTD and surgical donation programs.

This work has been supported in part by the Tissue Core, the Molecular Genomics Core Facility, the Proteomics and Metabolomics Core, the Biostatistics and Bioinformatics Shared Resource, the Analytical Microscopy Core, and the Bioimaging Nikon Center of Excellence at the H. Lee Moffitt Cancer Center & Research Institute, an NCI-designated Comprehensive Cancer Center (P30-CA076292). The contents of this work are solely the responsibility of the authors and do not necessarily represent the official views of the NCI or NIH.

## AUTHOR CONTRIBUTIONS

Conceptualization, S.A.A., H.D.A. and E.R.F.; methodology, S.A.A., H.D.A., V.Y.R., D.T.N. and W.G.S; investigation, S.A.A., H.D.A., V.Y.R., N.H., C.C., K.A.M., J.B., M.R., A.L., and A.Y.A.; analysis, S.A.A, H.D.A., V.Y.R., M.R., J.H.L., P.S., X.Y., A.Y.A, and J.M.K.; writing, S.A.A. and E.R.F.; review and editing, S.A.A., H.D.A., V.Y.R., J.H.L., and E.R.F.; visualization, S.A.A., H.D.A., V.Y.R., and X.Y.; resources, G.M.W., G.M.D., T.B., D.C., W.G.S., and E.B.H.; supervision, E.R.F.; funding acquisition, E.B.H. and E.R.F.; all authors reviewed and approved the manuscript.

## DECLARATION OF GENERATIVE AI AND AI-ASSISTED TECHNOLOGIES IN THE WRITING PROCESS

Grammarly (Enterprise) was used for copyediting support (e.g., grammar, clarity, and tone) under Moffitt Cancer Center’s BAA-governed environment, with generative AI features disabled. The tool provided only non-generative, in-line suggestions and did not create content or contribute to the authors’ intellectual work.

## STAR METHODS

### EXPERIMENTAL MODEL AND STUDY PARTICIPANT DETAILS

#### Animal maintenance and studies

*Kras^LSL-G12D/+ (^*^6)^ *(RRID:IMSR_JAX:008179)* and *Kras^LSL-G12D/+^;p53^fl/fl^ ^(^*^7)^ (RRID:IMSR_JAX:032435) mouse models were crossed with *Rosa^mTmG/mTmG^* ^(60)^ (RRID:IMSR_JAX:007676) to obtain *Kras^LSL-G12D/+^;Rosa^mTmG/mTmG^*and *Kras^LSL-G12D/+^;p53^fl/fl^;Rosa^mTmG/+^*mice on a C57BL/6 background. The *Kras^LSL-G12D/+^;Rosa^mTmG/mTmG^* mice were additionally crossed with *p53^R172H/+ (^*^7)^ (RRID:IMSR_JAX:008652) to generate *Kras^LSL-G12D/+^;p53^LSL-R172H/+^;Rosa^mTmG/mTmG^* mice. Furthermore, *Kras^LSL-G12D/+ (^*^6)^ *(RRID:IMSR_JAX:008179)* mice were crossed with a conditional knockout allele of *TAp73^fltd/fltd^* to generate *Kras^LSL-G12D/+^;TAp73^fltd/fltd^* ^(61)^ mice on a C57BL/6 background.

All studies were conducted in accordance with established protocols approved by the Institutional Animal Care and Use Committee (IACUC) of H. Lee Moffitt Cancer Center. Mice were housed in a 12h light/dark cycle, with room temperature at 21 ± 2 °C and humidity between 45 and 65% in pathogen-free and ventilated cages. They were allowed free access to irradiated food and autoclaved water. Both female and male mice, 6-8 weeks old, received 7.5 × 10^7^ PFU of an adenovirus expressing *Cre* recombinase under the CMV promoter (Ad5-CMV-Cre; Baylor College of Medicine) via intratracheal instillation to initiate tumor formation. Mice were monitored, and tumors were collected at the endpoint in accordance with IACUC guidelines. The endpoint criteria included large tumor burdens by magnetic resonance imaging (MRI; > 500 mm^3^), onset of dyspnea, hunching, or significant body weight loss (10% or more).

#### Patient tissue collection

Patient tissues were received through MCC20023 and the Rapid Tissue Donation (RTD) protocol^21^ at Moffitt Cancer Center (MCC17820 and MCC18843). All patients provided written informed consent, and the Moffitt Cancer Center Institutional Review Board approved the collection protocols. For the RTD program, when feasible, a discussion of the study and informed consent were provided during clinical care. Patient tissues were obtained during lung surgery or, for the RTD program, during an autopsy at Moffitt Cancer Center, in coordination with and with the permission of the Pathology Department Autopsy Director.

For both OR and RTD cases, primary lung tumors and metastatic tissues from liver or brain sites were collected and immediately placed on HPLM media with 1X Penicillin-Streptomycin (Thermo Scientific, 15140122). Tissues were kept on ice until processed for microtumor culture, histology, and cryopreservation.

#### Mouse tissue collection

For microtumor and organoid cultures, mice were euthanized with CO_2_ and cervical dislocation. In a sterile biosafety cabinet, the chest cavity was opened, and the heart and lungs were perfused with cold sterile PBS. The tumors were dissected from the lungs, guided by GFP in mice with the Rosa^mTmG/mTmG^ genotype or Tdtomato fluorescence in *Kras^LSL-G12D/+^;TAp73^fltd/fltd^*mice using fluorescence visualization goggles (BLS Ltd.). Multiple tumors were collected from each mouse’s lungs and pooled in 1X PBS on ice for omics, microtumor, and/or organoid processing.

#### Preparation of mouse and human microtumor culture

LUAD microtumors were cultured in HPLM media supplemented with 10% FBS (Sigma Aldrich, F0926-500), 1% Penicillin-Streptomycin (10,000 U/mL; Gibco, 15140122), 100 µM HEPES (Gibco, 15630-080), 1% ITS supplement media (Corning, 25-800-CR), and 0.5 µM A83-01 (Tocris, 2939). Additionally, all media for patient samples had 1:1000 Fungin (Invivogen, ant-fn-1). The day before plating the microtumors, an aliquot of the liquid-like solid microgels (LLS^TM^; Aurita BioScience) was centrifuged for 8 minutes at 1500 x *g*. The storage solution was replaced with HPLM media, and the LLS was equilibrated overnight at 37 °C.

Microtumors were prepared using the methodology described previously^19,20^, with minor changes specific to lung adenocarcinoma culture. Briefly, isolated tumors from mouse lungs or patient samples were minced using sterile surgical scissors until the tissues were approximately ≤ 1 mm^3^. The resulting microtumors were washed 3 times in 1X PBS to remove residual single cells, extracellular matrix, and blood. Then, they were suspended in the media-equilibrated LLS. A 200 µL LLS and microtumor mixture containing an average of 30 microtumors was added to each well of the 24-well Darcy Plate^TM^ (Aurita BioScience, DRCY24V). The plates were spun at 200 x *g* for 5 minutes using moderate acceleration and deceleration settings to settle the LLS and microtumors into the wells. After centrifugation, 5 mL of media was carefully added to each of the four quadrants of the Darcy plate, and a custom battery-powered air pump was connected to the vacuum port to create negative pressure in the bottom chamber and maintain constant perfusion at approximately 250 µL/hr. The plates were incubated at 37 °C and 5% CO_2_. The quadrants were refilled with fresh media every 24 hours, and the flow-through media was discarded. Microtumors were cultured in Darcy plates for 10 days before collection.

To harvest the microtumors after culture, they were transferred to 15 mL conical tubes using pre-wetted wide-bore pipette tips and washed three times with 1X PBS to remove LLS particles. Then, the microtumors were processed for downstream analysis, such as immunofluorescence, flow cytometry, and omics.

#### Preparation of organoid culture

Mouse lung tumors were minced as described above and then dissociated to single cells for 15 minutes at 37°C in 0.2% Collagenase I (Gibco, 17100-017); 0.2 % Dispase II (Roche, 04942078001); 25 U/mL DNase I (Sigma, D4527-10KU) in HPLM supplemented with 10 mM HEPES and 1% Pen-Strep. The samples were passed through a syringe fitted with an 18-gauge blunt-tip needle to dissociate the tissue, then neutralized with a 4-fold dilution in PBS containing 2% fetal bovine serum (FBS). Cells were strained using a 100 µm nylon mesh cell strainer (Falcon, 08-771-19) and centrifuged at 300 x *g* for 5 minutes. In ultra-low attachment 24-well plates (Corning, 3473), the dissociated cells were plated at a density of 100,000 cells per well in 50 µL domes of 100% Cultrex RGF Basement Membrane Extract, Type 2 (R&D Systems, 35-330-0502). The domes were incubated at 37 °C for 15 minutes to allow the Cultrex to solidify, then fresh media was added. Lung organoids were cultured in the same media recipe as the microtumors: HPLM media supplemented with 10% FBS, 1% pen-strep, 100 µM HEPES, 1% ITS supplement media, and 0.5 µM A83-01 and were incubated at 37 °C and 5% CO_2_. Organoids were cultured for 3-4 passages using mechanical dissociation, then collected for downstream analysis.

## METHOD DETAILS

### Microtumor Immunofluorescence

Microtumors were collected from the perfusion plates as described above. They were subsequently fixed in 10% neutral buffered formalin overnight at 4°C. The formaldehyde was washed out with 1X PBS, and the microtumors were then permeabilized for 1.5 hours in 0.5% Triton X-100 in PBS, followed by a 15-minute wash in 1X PBS. Samples were blocked in 3% BSA in PBS for 2 hours. Primary antibodies were diluted 1:50 in 3% BSA in PBS and incubated overnight at 4°C. For unconjugated primary antibodies, samples were washed for 15 minutes in 1X PBS, followed by an overnight incubation at 4°C with a secondary antibody at a 1:50 dilution in 3% BSA in 1X PBS. To visualize actin, phalloidin-iFluor 488 Reagent (Abcam, ab176753) was added at a 1:100 dilution to the final overnight antibody incubation. After antibody incubation, microtumors were washed for 15 minutes in 1X PBS and incubated for 30 minutes in Hoechst 34580 (1:500; BD Biosciences, 565877) in 1X PBS. Next, samples were washed for 15 minutes with 1X PBS and placed in glass-bottom 35 mm dishes (Cellvis; D35-20-0-N) for confocal imaging. The antibodies used were mouse CD45 monoclonal (clone D3F8Q) AF647-conjugated primary (Cell Signaling Technology, Cat# 81143, RRID: AB_2943239); human CD45 monoclonal (clone HI30) APC-conjugated primary (Thermo Fisher Scientific, Cat# 51-0459-42, RRID: AB_2802301); and Ki67 polyclonal primary (Abcam, Cat# ab15580, RRID: AB_443209).

### Live/Dead staining on microtumors

Microtumors were collected from the perfusion plates using pre-wetted wide-bore pipette tips and transferred to 1.5 mL microcentrifuge tubes. They were washed twice with fresh media and stained using the LIVE/DEAD™ Cell Imaging Kit (488/570; R37601; Invitrogen). Microtumors were stained following the manufacturer’s instructions, and the nuclei were stained using Hoechst 34580 (1:500; BD Biosciences, 565877). Microtumors were imaged immediately after dye incubation.

### Microtumor confocal imaging

Microtumor imaging datasets were acquired with a Nikon AXR confocal (Nikon Instruments, Inc., USA) mounted on a Nikon Ti2E inverted microscope and controlled by the NIS-Elements imaging software (Nikon Instruments, Inc., USA). Confocal images were collected with a 10X Plan APO Lambda D air objective (NA 0.45) or 20X Plan APO Lambda D air objective (NA 0.80). Z-stacks (1 µm step size, 1024×1024) were acquired and visualized using maximum-intensity projection.

### Flow cytometry

Single cell suspensions were generated from primary tumors or microtumors by digesting with 1 mg/mL Collagenase IV (Worthington, LS004188) and 25 U/mL DNase I (Sigma, D4527-10KU) in HBSS (Corning, 21-023-CV) for 30 minutes at 37°C. The samples were passed through a syringe fitted with an 18-gauge blunt-tip needle to dissociate the tissue, then neutralized by a 3-fold dilution in PBS containing 2% fetal bovine serum (FBS). Cells were strained using a 70 µm nylon mesh cell strainer (Falcon, 08-771-2) and centrifuged at 300 x *g* for 5 minutes. Single cell suspensions were generated from organoid cultures by first digesting for 10 minutes at 37°C with 2 mg/ml Dispase II (Roche, 04942078001) and 25 U/mL DNase I (Sigma, D4527-10KU) and then for 5 minutes at 37°C with 0.25% Trypsin/EDTA (Corning, 25053CI) and 25 U/mL DNase I. Cells were stained with a near-IR viability dye (Invitrogen, L34975) according to the manufacturer’s protocol. Cells were then treated with TruStain FcX PLUS (BioLegend, 156604) and stained on ice for 30 minutes in the dark with the following antibody cocktail diluted in BD Horizon Brilliant Stain Buffer (BD 563794): anti-CD45 BV711 (BD Biosciences, Cat# 563709, RRID: AB_2687455), anti-CD31 BV711 (BD Biosciences, Cat# 740680, RRID: AB_2740367), anti-Ep-CAM BV421 (BD Biosciences, Cat# 563214, RRID: AB_2738073), and anti-Sca1 AF647 (BD Biosciences, Cat# 565355, RRID: AB_2739205). Cells were then washed with FACS buffer (PBS + 1% BSA) and transferred to flow tubes through a 35 µm strainer cap. Data were acquired on a BD LSRII and analyzed with FlowJo version 10, according to the gating strategy described in Lee et al.^32^.

### Single-cell RNA sequencing sample preparation

Primary tumors were dissected from the mouse lungs as described above and minced. Microtumors were collected from the Darcy plates and washed 3 times with 1X PBS to remove the LLS particles. Samples were digested with 0.2% Collagenase I (Gibco 17100-017, diluted 1:50 from 100 mg/mL solution), 0.2% Dispase II (Roche 04942078001, diluted 1:5 from 10 mg/mL solution), and 25 U/mL DNase I (Sigma D4527-10KU, diluted 1:143 from 1 mg/mL solution) in MEM. After a 15-minute incubation at 37 °C in the digestion media, the samples were passed through a syringe with an 18-gauge blunt-tip needle to dissociate the tissue. 2% FBS in 1X PBS was added to the samples, which were then filtered through a 40-µm nylon mesh (Falcon, 352350). A Red Blood Cell (RBC) lysis (BioLegend, Cat #: 420301) step was performed following the manufacturer’s instructions. Organoids were dissociated to the single cell level as described above. Samples were resuspended in 100 µL of PBS + 0.4 mg/mL BSA, and cell count and viability were assessed using AO/PI staining (Nexcelom, ViaStain™ AOPI Staining Solution, CS2-0106-25mL).

### Single-cell RNA sequencing

Single-cell RNA sequencing was performed using the 10X Genomics Chromium system. A single-cell suspension derived from dissociated mouse tumors, microtumors, or organoids was analyzed for viability using the Nexcelom Cellometer K2 and then loaded onto the 10X Genomics Chromium Single Cell Controller at a concentration of one thousand cells per microliter to encapsulate approximately 6,000 cells per sample. Briefly, single cells, reagents, and 10x Genomics gel beads were encapsulated into individual nanoliter-sized gel beads in Emulsion (GEMs), and then reverse transcription of polyadenylated mRNA was performed within each droplet at 53°C. The cDNA libraries were then completed in a single bulk reaction using the 10X Genomics Chromium NextGEM Single Cell 3’ Reagent Kit v3.1, and 50,000 sequencing reads per cell were generated on the Illumina NovaSeq 6000 instrument. Demultiplexing, barcode processing, alignment, and gene counting were performed using the 10X Genomics CellRanger v7.1.0 software with the GRCm39-2024-A reference.

### Single-cell RNA-seq data processing, filtering, batch effect correction, and clustering

Raw sequencing reads from scRNA-seq were processed using Cell Ranger (v7.1.0, 10X Genomics). Briefly, the base call (BCL) files generated by Illumina sequencers were demultiplexed into fastq files based on the sequences of the sample index and aligned against GRCm38 mouse transcriptome using STAR^62^. Cell barcodes and UMIs associated with the aligned reads were corrected and filtered. Filtered gene-barcode matrices containing only barcodes with UMI counts that passed the cell-detection threshold were imported into Seurat v4.0.^63^ for downstream analysis. Barcodes with fewer than 200 genes expressed or more than 10% UMIs originating from mitochondrial genes were filtered out; genes expressed in fewer than 3 barcodes were also excluded. For each sample, standard library size and log-normalization were performed on raw UMI counts using *NormalizeData*, and the top 5000 most variable genes were identified by the “best” method in *FindVariableFeatures*.

Individual data were further integrated to remove batch effects using an anchor-based method^64^ implemented in Seurat v4.0 using *FindIntegrationAnchors* and *IntegrateData* functions in Seurat with 8,000 “anchors” and top 40 principal components. Briefly, dimension reduction was performed on each data set using diagonalized canonical correlation analysis (CCA), and L2-normalization was applied to the canonical correlation vectors to project the datasets into a shared space. The algorithms then searched for mutual nearest neighbors (MNS) across cells from different datasets to serve as “anchors”, which encoded the cellular relationship between datasets. Finally, correction vectors were calculated from “anchors” and used to integrate datasets.

From the integrated data, scaled z-scores for each gene were calculated using the *ScaleData* function in Seurat by regressing against the percentage of UMIs originating from mitochondrial genes, S and G2/M phase scores, and total reads count. A shared nearest neighbor (SNN) graph was constructed based on the first 40 principal components computed from the scaled integrated data. Louvain clustering was performed using the *FindClusters* function at resolution 1 for major cell-type scRNA-seq data (47 clusters). Uniform manifold approximation and projection (UMAP) was used to visualize single-cell gene expression profiles and clustering, using the *RunUMAP* function in Seurat with default settings. Differential expression analysis was performed using *FindAllMarkers* function in Seurat with logfc.threshold=0.25, min.pct=0.1, and test.use=”wilcox”. Cells within each cluster were compared against all other cells. Genes with Bonferroni-corrected p-value < 0.05 and an average log-fold change > 0.25 were considered differentially expressed. Clusters were annotated by comparing differential genes with canonical markers for major populations: B cells (*Cd79a, Cd79b, Cd19*), T cells (*Cd3e, Cd3d, Cd8a, Cd4*), NK cells (*Klrb1c, Ncr1, Nkg7*), Macrophages (*Cd68, Mrc1, C1qc, C1qb*), Monocytes (*Lyz2, Vcan, Chil3, Fn1*), Neutrophils (*S100a8, S100a9, Cxcl2*), cDC1 (*Xcl1, Cd36, Itgae*), cDC2 (*Itgax, H2-DMb1, Mgl2*), mregDC (*Fscn1, Ccl22, Cacnb3*), pDC (*Siglech, Ccr9, Bsl2*), Mast cells (*Mcpt8*, *Cd200r3*, *Ms4a2*), Epithelial cells (*Sftpc, Krt7, Krt18*), Endothelial cells (*Pecam1, Cdh5*), Fibroblasts (*Col1a1, Col1a2*).

### Annotation of epithelial cells

Epithelial cells were extracted for further clustering analysis with a resolution of 0.6, yielding 19 clusters. These 19 clusters were annotated as malignant cells based on the positive expression of tumor markers (*Krt7, Krt18, Krt19*). To characterize the malignant cells, we compared the gene expression of the 19 malignant clusters with cell identity signatures previously reported in KP transplant mice^33–35^ and KACs cell type signature^36^. Enrichment scores of these signatures were calculated for each malignant cell using the *AUCell* algorithm implemented in SCENIC^65^, and the scores were visualized in heatmaps.

### Reagents and chemicals for mass spectrometry

All solvents and chemicals are LC-MS grade unless stated otherwise. Ammonium hydroxide, ammonium acetate, ammonium carbonate, ammonium formate, urea, HEPES, sodium orthovanadate, sodium pyrophosphate tetrabasic decahydrate, β-glycerophosphate disodium salt hydrate, and formic acid were all obtained from Sigma Aldrich (St. Louis, MO). Water, methanol (MeOH), acetonitrile (ACN), and isopropyl alcohol (IPA) were obtained from VWR (Radnor, PA). Trifluoroacetic acid (TFA) was obtained from Fisher Scientific (Waltham, MA). The Metabolomics Quality Control (QC) kit, comprising 14 isotopically labeled metabolite standards, was obtained from Cambridge Isotope Laboratories (Tewksbury, MA) and used as an internal standard (IS) for untargeted metabolomics. The Splash Lipidomix Mass Spec Standard (Avanti Polar Lipids, Alabaster, AL) kit was used as the IS for untargeted lipidomics analyses.

### Sample preparation for multi-omic mass spectrometry profiling

For all extractions, samples and reagents were maintained on ice throughout the process. Frozen tissue samples, organoid cultures, and pooled microtumor samples had 2 µL of metabolomics IS, followed by 400 µL of cold 80% MeOH, previously chilled for 1 hour at-80° C, for protein precipitation. Samples were homogenized vigorously for 30 seconds at 4° C at the highest intensity (Bullet Blender, Next Advance, Troy, NY), then incubated at-20° C for 30 minutes. Sample mixtures were centrifuged at 18,800 × g (Microfuge 22R, Beckman Coulter) at 4° C for 10 minutes. After centrifugation, the liquid supernatant was transferred to a new microcentrifuge tube and dried by vacuum concentration at room temperature (SpeedVac, Thermo). Dried samples were stored at - 80 °C pending analysis.

The protein pellets were further extracted for lipid analysis by adding 3 µL of the lipid IS and 300 µL of 100% IPA, then vortexing thoroughly for 30 seconds. Extracts were incubated for 30 minutes at-20° C before centrifugation at 18,800 × g at 4° C for 10 minutes. The supernatant was transferred to a new microcentrifuge tube after centrifugation and dried under vacuum concentration at room temperature. Dried lipid samples were stored in-80° C pending analysis, and protein pellets were reserved for further processing.

After pulverization and metabolite/lipid extractions, the protein pellets were resuspended under sonication (Bioruptor) in denaturing aqueous lysis buffer, containing 8M urea, 20 mM HEPES (pH 8), 1 mM sodium orthovanadate, 2.5 mM sodium pyrophosphate, and 1 mM β-glycerophosphate. Bradford assays were performed to determine the protein concentration of each sample. The proteins were reduced with 4.5 mM dithiothreitol (DTT) and alkylated with 10 mM iodoacetamide (IAAm) prepared as aqueous 10X stock solutions. Trypsin digestion (enzyme-to-substrate ratio 1:20) was conducted overnight at room temperature; another aliquot of trypsin was added for an additional 2-hour digestion. To quench proteolysis, the tryptic peptides were acidified with aqueous 1% trifluoroacetic acid (TFA) and desalted using C18 Sep-Pak cartridges (Waters) according to the manufacturer’s instructions.

Proteolytic peptides from each sample were labeled with TMTPro reagent. Label incorporation was assessed by LC-MS/MS and spectral counting to verify ≥95% incorporation in each channel. The 16 samples were then pooled and lyophilized. After lyophilization, the peptides were re-dissolved in 400 µL of aqueous 20 mM ammonium formate (pH 10.0); bRPLC A solvent was aqueous 5 mM ammonium formate, 2% acetonitrile, pH 10.0, and bRPLC B solvent was aqueous 5 mM ammonium formate, 90% acetonitrile, pH 10.0. The basic pH-reversed-phase separation was performed on a BEH C18 XBridge column (4.6 mm ID × 100 mm length, packed with 3.5 µm particles and 130 Å pore size, Waters). The peptides were eluted using the following gradient: 5% B for 10 minutes, 5% - 15% B in 5 minutes, 15-40% B in 47 minutes, 40-100% B in 5 minutes, and 100% B held for 10 minutes, followed by re-equilibration at 1% B. The flow rate was 0.6 ml/min, and 96 fractions were collected (1 minute per well) and concatenated into 24 for protein expression. Vacuum centrifugation (Speedvac, Thermo) was used to dry the peptides.

Further details on the omics acquisition are available in the Document S1.

### Mass spectroscopy data processing and analysis

Detailed methods on the omics data processing are available in the Supplementary Information file.

Data analysis of all omics datasets was performed using MetaboAnalyst 6.0 ^66,67^ using one-factor statistical analysis. All missing values were replaced with 1/5^th^ of the limit of detection (LOD), and feature lists were further filtered using interquartile range filtering for those containing more than 5,000 features. Pathway analyses for metabolomics and lipidomics datasets were performed using relative betweenness centrality in the KEGG database.

For differentially expressed protein identification between microtumors and primary tumors, the fold change for each identified protein was calculated, and a standard t-test was performed, and proteins with a p-value < 0.05 were considered differentially expressed. Following differential analysis, pre-ranked gene set enrichment analysis (GSEA)^55,56^ was performed.

### Preparation of patient tumors and microtumors for histological analysis

All tissue samples were fixed overnight at 4 °C using 10% neutral buffered formalin. The primary tumors were paraffin-embedded by the Tissue Core at Moffitt Cancer Center following standard protocols.

Prior to paraffin-embedding, 10-15 microtumors were embedded in 100 µL warm Histogel^TM^ specimen processing gel (Epredia, HG-4000-012) using a 10 x 10 x 5 mm Tissue-Tek Cryomold (Sakura, Cat #: 4565). Once the histogel solidified, the microtumors were removed from the mold and paraffin-embedded by the Tissue Core.

The formalin-fixed paraffin-embedded (FFPE) tissue blocks were sectioned into 4-μm sections, and all slides were stored at 4 °C until immunofluorescence staining.

### Multiplex immunofluorescence of SCLC markers

Formalin-fixed paraffin-embedded slides of primary tumors and microtumors were baked for one hour at 65 °C, and subsequently dewaxed using xylene and rehydrated using a series of ethanol dilutions. The slides were stained using the Opal Polaris 7-Color Manual IHC kit (Akoya Biosciences, NEL861001KT) according to the manufacturer’s instructions. The antibodies used were as follows: anti-ASCL1 (MASH1, Abcam, ab211327, 1:300, RRID: AB_2924270), anti-CD45 (Abcam, ab281586, 1:2000, RRID: AB_3697165), anti-NEUROD1 (Abcam, ab213725, 1:300, RRID: AB_2801303), anti-NCAM1 (CD56, Sigma, AB5032, 1:500, RRID: AB_11213653), anti-POU2F3 (Sigma, HPA019652, 1:200, RRID: AB_1855585), anti-YAP1 (Santa Cruz Biotechnology, sc101199, 1:100, RRID: AB_1131430). Slides were imaged with the PhenoImager Fusion (Akoya Biosciences) at 20X, 0.5um/pixel resolution. Multispectral Imaging Technology was used to reduce autofluorescence and fluorophore crosstalk.

### RNA and DNA isolation of patient samples

Minced primary patient tumors and microtumors were snap-frozen until RNA and DNA isolation. Frozen tumors were suspended in RTL Plus Lysis buffer with β-mercaptoethanol (β-ME) and homogenized using the Bullet Blender Tissue Homogenizer (Next Advance). Once homogenized, RNA and DNA were isolated from the tissues using the Qiagen All-Prep DNA/RNA Mini kit (80204) following the manufacturer’s instructions.

### qPCR of patient samples

Total RNA was isolated from patient samples as described above. Then, cDNA was synthesized using the qScript® Ultra SuperMix (QuantaBio, 95217) following the manufacturer’s protocol. Triplicate qPCR reactions were performed using the TaqMan Universal PCR Master Mix, No AmpErase UNG (Applied Biosystems; 4324018) and TaqMan probes for the genes of interest. Each reaction contained 20 ng cDNA per well and was run using the QuantStudio 6 Flex Real-Time PCR System. The samples were run at 50 °C for 2 minutes, 95 °C for 10 minutes, and 50 cycles at 95 °C for 15 seconds, then 60 °C for 1 minute. The TaqMan probes used were: CDH1 (Hs01023895_m1); MUC1 (Hs00159357_m1); AXL (Hs01064444_m1); VIM (Hs00958111_m1); and POL2RA (Hs00172187_m1).

### Moffitt STAR solid tumor assay

RNA and DNA samples were analyzed using the Moffitt STAR 2.0 panel, which is designed to analyze and interpret somatically altered gene sequences in human malignancies. This assay is derived from the original Moffitt Star panel^43^ and analyzes the coding DNA exons from 517 cancer-related genes, copy number alterations for at least 49 genes, and select gene fusions with known and novel partners and splice site variants from 55 genes. Next-generation sequencing (NGS) of the samples was conducted using the TSO500 assay (Illumina, Inc.) on the NovoSeq (Illumina, Inc.) platform. The procedure followed the Illumina Trusight Oncology 500 Library Prep Reference Guide^59^.

### RNA-STEP assay

A custom RNA expression panel consisting of 204 genes was developed at Moffitt Cancer Center using the NanoString platform. The RNA Salah Targeted Expression Panel (RNA STEP^42^) was designed to analyze the expression of therapeutic target or biomarker genes relevant to future clinical trials. Each run included a reference universal mRNA control from pooled human normal tissues (BioChain Institute, Inc., Newark, CA, Catalog number R4234565) to maintain batch consistency and normalize gene signals. Raw data files were processed and normalized with geNorm in NanoString nSolver 4.0 advanced analysis software. Normalized log2 ratios were calculated by subtracting the normalized log2 counts of the universal mRNA control genes from those of the individual samples.

#### ND-646 treatment

For patient samples treated with an ACC1/2 inhibitor, ND-646, the media was removed after 6 days of perfusion culture. Fresh media containing a DMSO vehicle or 4 µM ND-646 diluted from a 10 mM stock solution (SelleckChem, S8377) was added to the Darcy plates, and the media was refreshed daily. Microtumors were collected after 96 hours of treatment and processed for immunofluorescence and untargeted lipidomics analysis as described above.

## SUPPLEMENTAL INFORMATION

Document S1. Figure S1 and supplemental methods.

Table S1. Clinical information of OR patients, related to Table 1.

Table S2. Clinical information of RTD patients, related to Tables 2 and 3.

