## Supplementary material for "*Ex Vivo* Culture of Patient-Derived Primary, Metastatic, and Post-Mortem Lung Cancer Reveals Targetable States of EMT and Metabolic Plasticity": Document S1

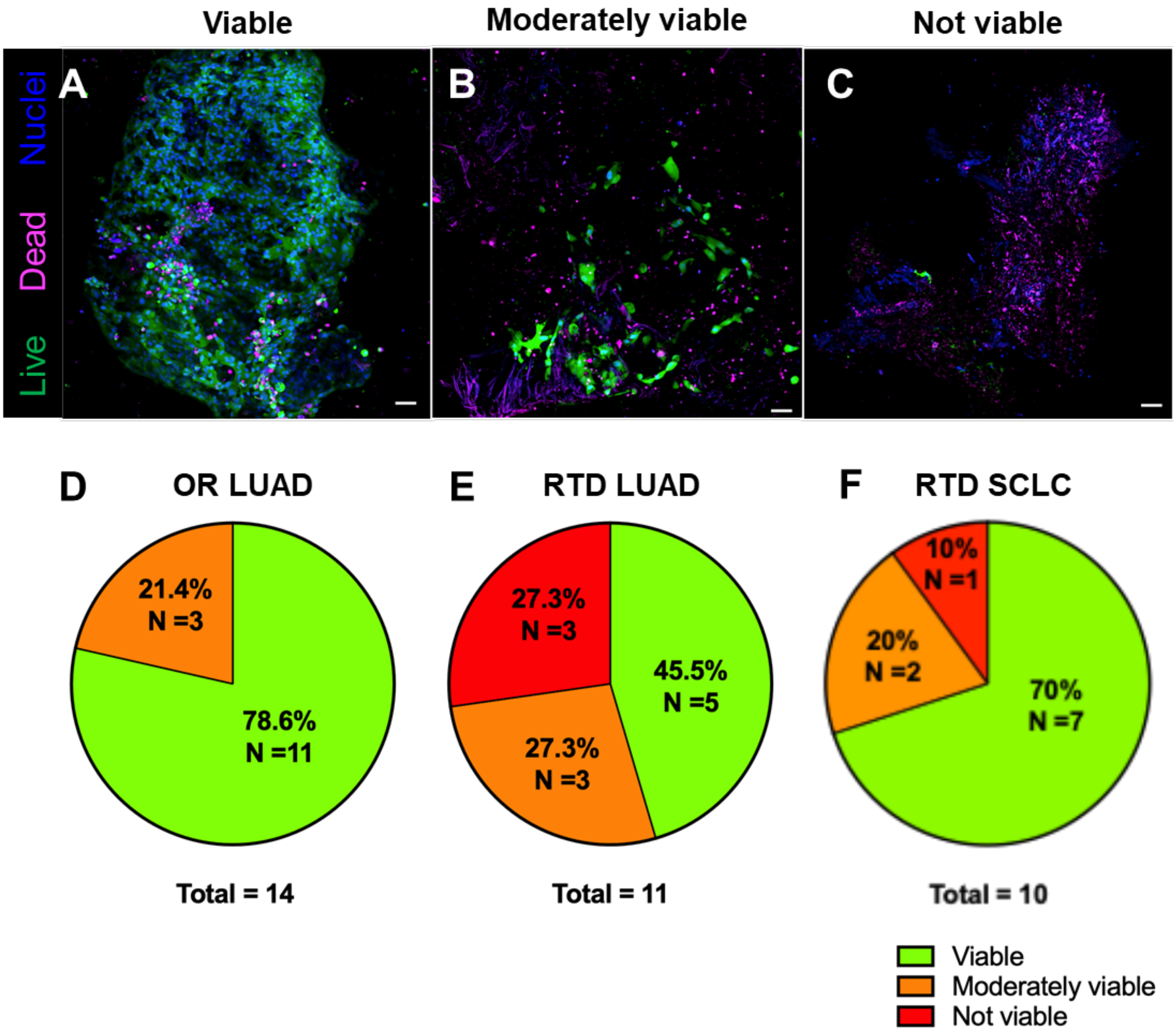


**Figure S1. LUAD and SCLC microtumors from patient OR and rapid tissue donation (RTD) sources are viable *ex vivo*. A – C:** Representative images showing viable, moderately viable, and non-viable microtumors after 10 days of perfusion culture. Live cells are shown in green, dead cells are in magenta, and all nuclei are in blue. Scale bar = 50 µm. **D – F:** Pie charts showing the percentage of viable, moderately viable, and non-viable patient tumors cultured *ex vivo* for OR LUAD samples, RTD LUAD samples, and RTD SCLC samples. The total sample count includes both the number of unique patients and the number of distinct tumor sites per patient.

### SUPPLEMENTAL METHODS

#### *Multi-omics mass spectroscopy data acquisition*

A nanoflow ultra high-performance liquid chromatograph (RSLC, Thermo, San Jose, CA) coupled to an electrospray bench top orbitrap mass spectrometer with high field asymmetric waveform ion mobility spectrometry for charge state selection (Orbitrap Exploris 480 with FAIMS, Thermo, San Jose, CA) was used for liquid chromatography tandem mass spectrometry peptide sequencing experiments. The sample was first loaded onto a pre-column (C18 PepMap100, 100 µm ID x 2 cm, 5 µm particles, 100Å pores) and washed for 8 minutes with aqueous 2% acetonitrile and 0.1% formic acid (solvent A). The trapped peptides were eluted onto the analytical column, (C18 PepMap100, 75 µm ID x 25 cm, 2 µm particles, 100 Å pores, Thermo, San Jose, CA). The 120-minute gradient was programmed as: 95% solvent A (aqueous 2% acetonitrile + 0.1% formic acid) for 8 minutes, solvent B (90% acetonitrile + 0.1% formic acid) from 5% to 38.5% in 90 minutes, then solvent B from 50% to 90% B in 7 minutes and held at 90% for 5 minutes, followed by solvent B from 90% to 5% in 1 minute and re-equilibrated at 5% B for 10 minutes. The flow rate on the analytical column was 300 nl/min. Cycle time was set at 1.5 seconds for data-dependent acquisition using two FAIMS compensation voltage values (-45 V and -65 V). Spray voltage was 2,100 V, and capillary temperature was 300 °C. The resolution values for MS and MS/MS scans were set at 120,000 and 45,000, respectively. Dynamic exclusion was set to 15 seconds for previously sampled peptide peaks.

Untargeted metabolomics was performed on reconstituted samples normalized by protein with 80% MeOH. Acquisition was performed using ultra high-pressure liquid chromatography (UHPLC) using a Thermo Scientific Vanquish interfaced with a Q Exactive HF high-resolution mass spectrometer (HRMS)(Thermo Scientific, San Jose, CA). Chromatographic separation was achieved on an Atlantis Premier BEH Z-HILIC VanGuard FIT column (2.1 mm ID x 150 mm length, 2.5 µm particle size) (Waters, Milford, MA). Gradient conditions were adapted from published methods[1]. Briefly, mobile phase A was aqueous 10 mM ammonium carbonate with 0.05% ammonium hydroxide in water, and mobile phase B was 100% ACN. The chromatography program began at 80% B with a linear decrease over 13 minutes until it achieved 20% B. It was isocratic at 20% B for 2 minutes, before returning to the original conditions within 0.1 minutes, where equilibration was performed for an additional 4.9 minutes. The total chromatography program was 20 minutes. The column was maintained at 30° C with a flow rate of 250 µL/min. Samples were acquired in positive and negative polarity with a 2 µL injection volume for each. Mass spectrometry was performed using heated electrospray ionization (HESI) operating at a spray voltage of 3.5 kV and 3.0 kV for positive and negative mode, a capillary temperature of 325° C, sheath gas and auxiliary gas of 50 and 10 arbitrary units (au), respectively. The mass range was 65 – 900 *m/z* in positive and negative ionization mode with a mass resolution of 120,000 using full scan acquisition.

Lipidomic samples were normalized to the protein concentrations and rehydrated with IPA as needed. Data acquisition was performed using UHPLC with tandem MS (LC-MS/MS) on the same system as the metabolomics analysis. Chromatographic separation was performed on a Brownlee SPP C18 column (2.1 mm x 75mm, 2.7 µm particle size, Perkin Elmer, Waltham, MA) using mobile phase A, which was water containing 0.1% formic acid and 1% of 1M ammonium acetate and mobile phase B, which was 1:1 ACN: IPA with 0.1% formic acid and 1% of 1M ammonium acetate. Gradient separation began with column equilibration for 2 minutes at 35% B, then a linear increase to 80% B over 6 minutes. A second linear increase to 99% B over 14 minutes was performed and maintained for 0.1 minutes before returning to equilibrium at 35% B for 3.9 minutes. The flow rate was maintained at 400 µL/min with a column temperature of 50°C. The lipid profile was monitored using data-dependent tandem mass spectrometry for the 5 highest-intensity ion signals (top 5) in positive and negative mode in separate experiments. Mass spectrometry was performed with HESI operating with sheath gas of 50 au, auxiliary gas 10 of au, and spray voltage of 3.5 kV. The capillary temperature was maintained at 325° C. The mass range for the acquisition was 120-1000 *m/z* with a mass resolution of 120,000 for MS and 30,000 for MS/MS in positive and negative ion modes. Isolation window width of 1.2 *m/z* was used for MS/MS with a stepped normalized collision energy (NCE) of 20, 30, and 40 au with a dynamic exclusion of previously sampled peaks for 8 seconds.

***Multi-omics mass spectroscopy data processing***

Metabolomics data were converted to open-source mzXML files using Raw Converter (Scripps) and then imported into MZmine 3.53 [2,3] for feature finding and alignment. Mass detection was set to a 10 ppm window with a 0.25 min RT tolerance. Metabolite identification was performed using an in-house RT standard library. Aligned and annotated data were exported using peak height values. For each positive and negative ion mode dataset, rows with fewer than 2 high-quality pre-gap-filled features were removed as low quality. Each dataset was then normalized separately with iterative rank order normalization, IRON[4], excluding heavy-labeled and unidentified rows from median sample identification (findmedian --pearson --iron-exclusions=unidentified_row_ids.txt --iron-spikeins=heavy_row_ids.txt) and normalization training (iron_generic --proteomics --iron-exclusions=unidentified_row_ids.txt --iron-spikeins=heavy_row_ids.txt). Normalized positive and negative ion mode datasets were then merged and $\log_2$ transformed, treating zero abundances as missing data. Each row was then annotated, through fuzzy compound name matching, against our internal manually curated HMDB, KEGG, and PubChem identifier table.

Peak picking and alignment of lipidomics data were performed using LipidSearch 4.2.2.1 [5,6] using the HCD targeted database with a QEX (Q Exactive) product search of 6 ppm for the precursor and product mass tolerances, a 1.0% MS/MS intensity threshold, and an m-Score threshold of 2. Quantitation mass tolerance of 6 ppm and a RT tolerance of ± 0.5 min was used. Alignment of the peak lists was performed using an LC-MS product search of the mean with an RT tolerance of 0.2 min and an m-Score threshold of 5.0. The aligned peak list was further filtered based on the most commonly observed adduct(s) for each lipid class. The filtered and aligned feature list was exported using peak heights and normalized using the same IRON pipeline described above for metabolomics datasets.

MaxQuant (version 1.6.14.0)[7] was used to identify peptides via the Andromeda search engine and quantify the TMT reporter ion intensities. Proteins were searched against murine entries in the UniProt database downloaded in April 2023. Up to 2 missed trypsin cleavages were allowed per peptide. Carbamidomethyl cysteine was set as a fixed modification, and methionine oxidation was set as a variable modification. Both peptide spectral match (PSM) and protein false discovery rate (FDR) were set at 0.01 using reverse sequences for the decoy. The match between runs feature was activated to enable identification from each LC-MS/MS run to be carried to other runs. Similar database searches were performed in Mascot (Matrix Science) to support data upload to PRIDE/ProteomeXchange[8,9]
